# Cognitive experience alters cortical involvement in navigation decisions

**DOI:** 10.1101/2021.12.10.472106

**Authors:** Charlotte Arlt, Roberto Barroso-Luque, Shinichiro Kira, Carissa A. Bruno, Ningjing Xia, Selmaan N. Chettih, Sofia Soares, Noah L. Pettit, Christopher D. Harvey

## Abstract

The neural correlates of decision-making have been investigated extensively, and recent work aims to identify under what conditions cortex is actually necessary for making accurate decisions. We discovered that mice with distinct cognitive experiences, beyond sensory and motor learning, use different cortical areas and neural activity patterns to solve the same task, revealing past learning as a critical determinant of whether cortex is necessary for decision-making. We used optogenetics and calcium imaging to study the necessity and neural activity of multiple cortical areas in mice with different training histories. Posterior parietal cortex and retrosplenial cortex were mostly dispensable for accurate decision-making in mice performing a simple navigation-based decision task. In contrast, these areas were essential for the same simple task when mice were previously trained on complex tasks with delay periods or association switches. Multi-area calcium imaging showed that, in mice with complex-task experience, single-neuron activity had higher selectivity and neuron-neuron correlations were weaker, leading to codes with higher task information. Therefore, past experience sets the landscape for how future tasks are solved by the brain and is a key factor in determining whether cortical areas have a causal role in decision-making.

## Introduction

Correlations between neural activity and decision-making have been studied extensively in the mammalian cortex, but the factors that determine whether cortical areas are actually necessary for decision tasks are not fully understood (Gold & Shadlen, 2007; Hanks et al., 2006; Katz et al., 2016; Salzman et al., 1990). Across studies, the necessity of cortical areas has been tested during a variety of decision-making tasks that involve different sensory, behavioral, and cognitive features (Ceballo et al., 2019; Erlich et al., 2011; Fischer et al., 2020; Goard et al., 2016; Guo et al., 2014; Harvey et al., 2012; Inagaki et al., 2018; Licata et al., 2017; Raposo et al., 2014; Yang & Zador, 2012; Zhou & Freedman, 2019; Znamenskiy & Zador, 2013). Collectively, these studies have formed proposals on the types of decisions for which specific cortical areas are essential. For example, in rodents, posterior parietal cortex (PPC) is necessary for visual, but not auditory, discrimination tasks (Licata et al., 2017) and is considered to be especially involved in tasks that have a short-term memory component, such as a delay period between sensory cues and choice reports, or a requirement for evidence accumulation over time (Lyamzin & Benucci, 2019). As another example, area LIP in monkeys is thought to be essential for sensory but not motor aspects of visual motion discrimination tasks (Zhou & Freedman, 2019). Notably, these studies have focused on how specific features of a task-of-interest determine which cortical areas are causally involved. However, in addition to the task-of-interest in a study, individual animals have learned a variety of associations throughout their lifetime and may have performed a diversity of tasks previously, often with different experiences between individuals. Although it is intuitive that past learning, beyond the demands of the task-of-interest, may impact how individuals make decisions, most studies of decision-making have not investigated the effect of past learning on the involvement of cortex. It therefore is not well understood how learning of previous tasks affects the necessity of cortical activity for decision-making.

Two previous studies investigated this topic in the sensory domain, comparing the involvement of visual area MT in depth and motion perception in monkeys with different perceptual experience (Chowdhury & DeAngelis, 2008; Liu & Pack, 2017). For coarse depth discrimination, MT was only necessary in monkeys that had no prior experience in fine depth discrimination tasks (Chowdhury & DeAngelis, 2008), showing a decrease in cortical involvement with additional experience. In contrast, for motion discrimination, previous training on moving dot stimuli rendered MT necessary for discriminating the motion of gratings (Liu & Pack, 2017), showing that sensory experience can also increase cortical involvement. Relatedly, studies of motor learning have shown that cortex is essential during the learning process but becomes dispensable after learning is completed (Hwang et al., 2019; Kawai et al., 2015). In these cases, the animal’s prior training was largely based on sensory or motor experience. However, studies have not investigated the impact of “cognitive experience”, which we broadly define as learning that extends beyond sensory or motor learning and includes learning of task rules and associations.

Here, we developed a paradigm to study the effects of previous task learning on the necessity and activity patterns of cortical areas. Mice performed a simple decision-making task in virtual reality, and we compared different groups of mice that either had or had not been previously trained on complex decision-making tasks. We used optogenetics and calcium imaging to measure the necessity and neural activity patterns of cortical areas during this simple task. Critically, we kept the sensory and movement aspects of the complex and simple tasks as identical as possible to test the effect of “cognitive experience” instead of perceptual or motor learning. We focused on areas of cortex that are thought to be critical for decision-making during navigation, including PPC, which converts sensory cues into motor plans (Freedman & Ibos, 2018), and retrosplenial cortex (RSC), which is critical for planning navigation trajectories (Alexander & Nitz, 2015).

We discovered that mice with different previous task experience used distinct sets of brain areas to solve the same simple task. During the simple decision task, mice without prior training on complex decision tasks performed well above chance levels when RSC or PPC was inhibited. In contrast, during the same simple task, mice with prior complex task training performed close to chance levels when these areas were inhibited. In addition, calcium imaging revealed that prior complex task experience resulted in increased selectivity of neural activity patterns for task-relevant variables and decreased correlations in neural activity during the simple task. Thus, individuals with distinct cognitive experience make outwardly identical decisions using different combinations of brain areas and neural activity patterns. We suggest that, because neural circuits are optimized for a wide range of computations beyond the ones required by a current task-of-interest, a global set of constraints and optimizations can dramatically impact the cortical areas that are necessary for decisions.

## Results

### Increased necessity of cortical association areas in complex versus simple decision tasks

We developed a paradigm in which head-restrained mice running on a spherical treadmill were trained to use visual cues to make navigation decisions in a virtual reality Y-maze (Figure 1A-B). We used this paradigm to create a “simple task” and two “complex tasks”. In the simple task, mice learned to associate visual cues – horizontal and vertical bars – with left and right turns, respectively, to obtain rewards. In this task, the visual cue was present throughout the entire Y-maze, and the rewarded cue-choice associations (e.g., horizontal bars-left choice) did not change (Figure 1C).

**Figure 1.**
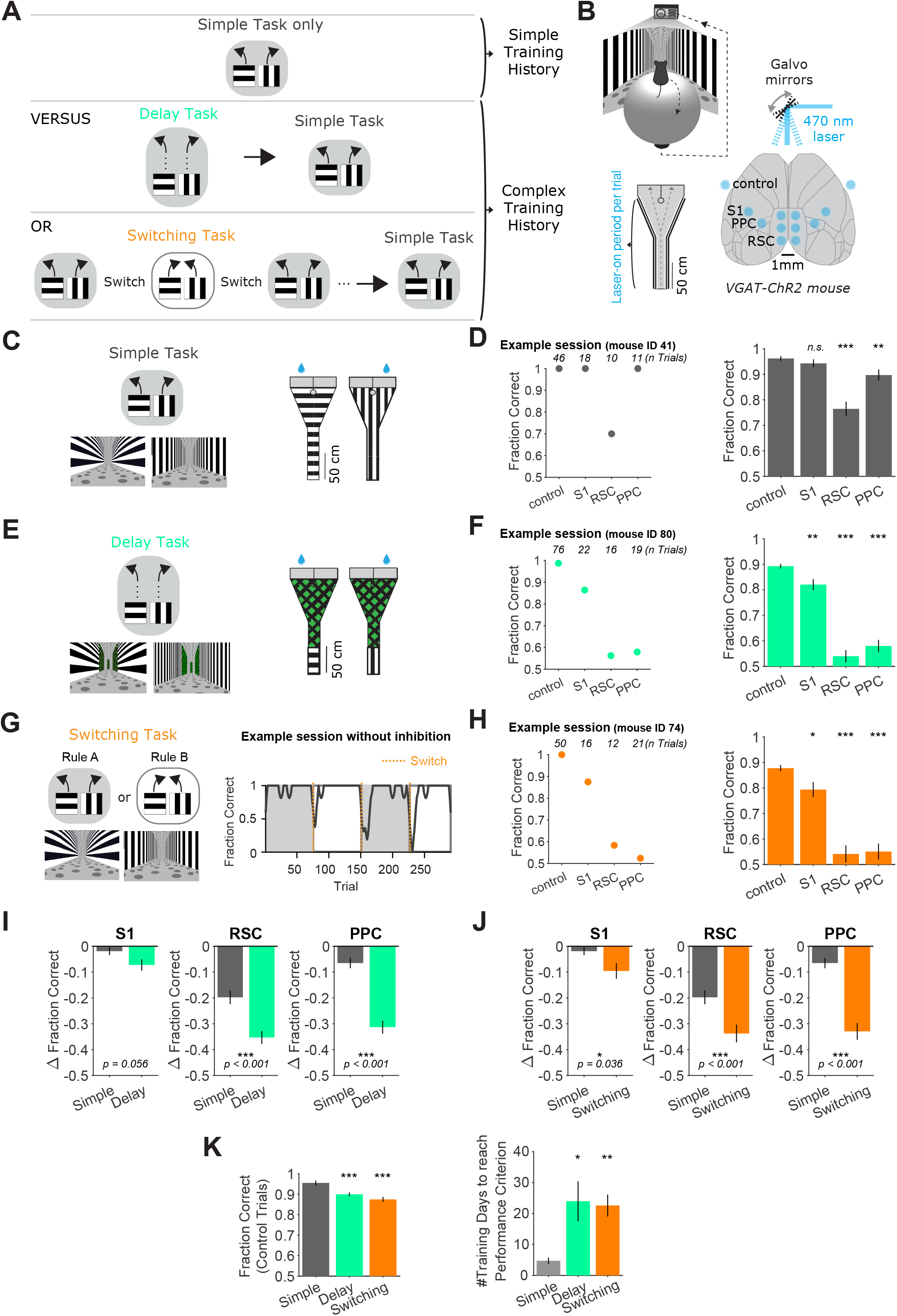
Increased necessity of cortical association areas in complex versus simple decision tasks. **(A)** Schematic overview of the behavioral tasks and task training sequences used in this study. Top row: One group of mice is trained in the simple task only. Middle row: Another group of mice is trained on the delay task and then transitioned to the simple task. Bottom row: Another group of mice is trained on the switching task and then transitioned to the simple task. The middle and bottom rows indicate complex training histories. **(B)** Top: Schematic of virtual reality behavioral setup. Bottom right: Schematic of optogenetic inhibition with bilateral target locations. Bottom left: Top view of Y-maze. Inhibition lasted from trial onset throughout maze traversal. **(C)** Left: Simple task schematic indicating two trial types (horizontal or vertical cues) and corresponding rewarded navigation decisions (running left or right). Corresponding VR screenshots at the trial start are below. Right: Top view of the two maze schematics. Water drops indicate hidden reward locations. **(D)** Left: Example session in the simple task showing mean performance for each inhibited location. Right: Performance in the simple task for each inhibited location across 45 sessions from 4 mice. Bars indicate mean ± sem of a bootstrap distribution of the mean. S1 p = 0.84; RSC p < 0.001; PPC p = 0.006; from bootstrapped distributions of ΔFraction Correct (difference from control performance) compared to 0, two-tailed test, α = 0.05 plus Bonferroni correction. Sessions per mouse: 11 ± 2. Trials per session: 53 ± 23 (control), 19 ± 8 (S1), 18 ± 9 (RSC), 20 ± 9 (PPC), mean ± SD. **(E)** Similar to **(C)**, but for the delay task. **(F)** Similar to **(D)**, but for the delay task. 62 sessions from 7 mice. S1 p = 0.006; RSC p < 0.001; PPC p < 0.001. Sessions per mouse: 9 ± 4. Trials per session: 60 ± 15 (control), 16 ± 6 (S1), 15 ± 4 (RSC), 17 ± 5 (PPC), mean ± SD. **(G)** Left: Schematic of the switching task, utilizing the identical mazes as the simple task. The cue-choice associations from the simple task (Rule A) were switched within a session (to Rule B). Right: Behavioral performance from an example session. Dotted orange lines indicate rule switches. **(H)** Similar to **(D)**, but for the switching task, Rule A trials only. 89 sessions from 6 mice. S1 p = 0.036; RSC p < 0.001; PPC p < 0.001. Sessions per mouse: 15 ± 5. Trials per session: 26 ± 9 (control), 8 ± 3 (S1), 7 ± 4 (RSC), 8 ± 3 (PPC), mean ± SD. **(I)** Comparison of inhibition effects (ΔFraction Correct) in the simple and the delay tasks for each cortical inhibition location. Bars indicate mean ± sem of a bootstrap distribution of the mean; two-tailed comparisons of bootstrapped ΔFraction Correct distributions, α = 0.05. Same datasets as in **(F, G)**. **(J)** Similar to **(I)**, but for the simple vs. switching task (Rule A trials only). Same datasets as in **(F, H)**. **(K)** Left: Comparison of performance on control trials across tasks, using only the first two laser-on blocks in each session. Bars indicate mean ± sem of a bootstrap distribution of the mean. Delay vs. simple p < 0.001; switching vs. simple p < 0.001; two-tailed comparisons of bootstrapped Fraction Correct distributions, α = 0.05. Right: Number of training sessions needed to reach performance criteria across tasks (Methods). Bars indicate mean ± sem across mice, n = 4 for simple task, n = 5 for delay task, n = 6 for switching task. Both delay and switching task data were compared to the simple task data using an unpaired two-sided t-test. Delay vs. simple p = 0.04; switching vs. simple p = 0.006.

The complex tasks were designed based on the same Y-maze concept and used the identical horizontal and vertical bars as visual cues. In the “delay task”, the visual cues were only present at the beginning of the maze, followed by a neutral visual pattern on the walls for the remainder of the maze (Figure 1E). This neutral pattern was identical across trials and did not provide information about the reward location. This design was based on the commonly used approach of inserting a delay period between the sensory cues and choice reports and has been used previously in navigation decision tasks (Driscoll et al., 2017; Harvey et al., 2012). In the “switching task”, the rewarded relationships between the visual cues and left-right choices were switched across blocks within a single session, resulting in two rules (Rules A and B) (Figure 1G). The same visual cue was thus associated with left choices in one rule block and right choices in the other rule block. The current rule and rule switch were not explicitly signaled, so the mouse learned rule switches by trial and error and stored a belief of the current rule in memory. Therefore, after rule switches, a mouse’s performance dropped and then recovered to high and stable levels after tens of trials (Figure 1G). For one of the two rules (Rule A), the trials were the same as in the simple task, including the same cues and rewarded choices. In fact, the software code used to create the virtual environments was identical between the simple task and Rule A of the switching task. Across both the simple and complex tasks, the mice experienced the same choice-informative sensory cues and had to run through similar or identical mazes to report their choices. Thus, the key difference between the simple and complex tasks was due to “cognitive” complexity through the addition of a delay period or frequent switches in the associations, rather than differences in the sensory cues informing choices or differences in motor output.

We tested the necessity of various cortical areas during the simple and complex tasks using optogenetics to activate GABAergic interneurons, which leads to silencing of nearby excitatory neurons (Guo et al., 2014; Li et al., 2019; Minderer et al., 2019). Channelrhodopsin-2 (ChR2) was expressed in inhibitory interneurons in transgenic mice and photostimulated using a clear-skull preparation with a two-dimensional laser scanning system (Figure 1B). We focused inhibition on two cortical association areas previously linked to decision-making and navigation, PPC and RSC (Driscoll et al., 2017; Fischer et al., 2020; Harvey et al., 2012). To match each area’s anatomical extent, we used three inhibition spots in each hemisphere for RSC and one spot per hemisphere for PPC. As a control, we inhibited a spot in primary somatosensory cortex (S1), an area not implicated in visual decision-making (Figure 1B). Inhibition trials were interleaved with control trials in which the laser spot was positioned outside the mouse’s brain. On inhibition trials, photostimulation was applied bilaterally and throughout the duration of the mouse’s maze traversal.

We first considered inhibition effects on performance of the simple task in mice that had only been trained in the simple task (Figure 1D). Silencing S1 did not affect performance, and inhibition of PPC resulted in a very small performance decrease, with mean performance of 90 ± 2% correct (mean ± SEM). Inhibition of RSC had the largest effect, resulting in intermediate performance levels of 77 ± 3% correct. However, mice still performed well above chance (50% correct). To assess whether the effect of RSC inhibition was specific to the cognitive requirements of the simple task or related to lower-level processes required for task performance such as vision, movement, and basic navigation, we silenced the same areas in an even simpler task in which mice ran towards a visual target present on either side of the maze end to obtain rewards (Harvey et al., 2012). Effects on performance were similar in this run-to-target task, suggesting that RSC inhibition in the simple task may impair lower-level processes such as basic navigation instead of decision-making based on cue-choice associations (Figure 1—figure supplement 1). Together, these results indicate that the cortical areas we silenced were only modestly involved in the decision in the simple task.

We next considered the effects of inhibition on mice performing the delay task or the switching task (Figure 1,E,F,G,H). In the switching task, we silenced cortical areas during the periods of high performance after accuracy had recovered following a rule switch (Figure 1G). S1 inhibition during the complex tasks caused a modest decrease in performance, but mice still performed the tasks at high levels (Figure 1F, H). In contrast, inhibition of PPC or RSC greatly impaired performance in the delay and switching tasks and resulted in performance of 55 ± 3% correct, which is close to chance levels. With PPC and RSC inhibition, many mice exhibited biases in the choices they made, whereas others appeared to choose randomly between left and right (Figure 1 —figure supplement 2). The effects of inhibiting PPC and RSC were markedly larger in the delay and switching tasks than in the simple task (Figure 1I-J). Therefore, adding a delay epoch to the trial or frequent association switches across trials increased the necessity of cortex relative to the simple task.

We also asked whether the increased cortical necessity was especially apparent in the task period that was changed compared to the simple task. In the delay task, we thus restricted photoinhibition to the cue or the delay segment of the trial (Figure 1—figure supplement 3). Inhibition of PPC or RSC in both trial segments decreased performance to a greater degree than in the simple task, indicating that PPC and RSC are necessary for multiple epochs of the task. In the switching task, the main difference relative to the simple task is the introduction of association switches, so activity in PPC and RSC may be necessary for storing or updating the rule. Taking advantage of interleaved control and inhibition trials, we found that the effects of inhibition were restricted to the current trial, as inhibition did not affect subsequent control trials (Figure 1—figure supplement 4). We also looked for a role in updating the rule, which is likely critical following the completion of a trial, in particular after a rule switch. We focused on PPC as a candidate for updating the rule because PPC has sensory- and choice-related history signals (Akrami et al., 2018; Morcos & Harvey, 2016). We silenced PPC during the inter-trial interval on every trial following a rule switch, for 50 consecutive trials. However, PPC inhibition did not affect the recovery of performance after a rule switch (Figure 1—figure supplement 4). Together, these results suggest that the large inhibition effects during the switching task are not due to impaired storing or updating of the rule. Thus, the increased necessity of PPC and RSC during the delay and switching tasks did not appear to be specific to the added task components.

Overall, our results indicate that more complex tasks require cortical activity, specifically in PPC and RSC, to a larger extent than a simple task. We verified that the complex tasks were indeed more challenging for mice. Relative to the simple task, it took mice longer to become experts at the delay and switching tasks (see Methods), and their performance on control trials was lower (Figure 1K). These findings are consistent with, and extend, previous work that concluded cortical necessity increases with task complexity (Harvey et al., 2012; Pinto et al., 2019).

### The necessity of PPC and RSC depends on a mouse’s previous cognitive experience

This set of tasks provided a platform for testing the effects of prior learning on cortical necessity for decision-making in the simple task, by comparing groups of mice with or without previous training on the complex tasks. A first group of mice was only trained on the simple task. The second group was first trained on one of the complex tasks and then transitioned to the simple task for 14 consecutive sessions (one session per day), without experiencing the complex task again. Different cohorts of mice were transitioned to the simple task from the switching task and delay task. This design allowed us to compare different mice performing the same task (the simple task) but with distinct training histories.

We first considered the mice that were trained to be experts on the delay task and then transitioned to the simple task (Figure 2A). In these mice, inhibition of PPC and RSC during the simple task greatly impaired behavioral performance to mean levels of 67 ± 3% and 62 ± 2% correct (mean ± SEM), respectively (Figure 2B-C). Thus, these mice needed PPC and RSC activity to perform at high levels on the simple task. These results were strikingly different from the inhibition effects in mice trained only on the simple task. Without complex-task experience, mice performed at 90 ± 2% and 77 ± 3% correct with PPC or RSC inhibited (Figure 2C), respectively, indicating they did not strongly rely on PPC or RSC activity for task performance. The larger effect of PPC and RSC inhibition due to previous delay task experience was not only apparent immediately after the transition to the simple task but persisted for the full two weeks that we investigated (Figure 2D-E). Therefore, the effect of PPC and RSC inhibition had markedly larger effects in the simple task when mice had previous training in the delay task, both in the first and second week after the task transition. This persistent difference is particularly surprising because the task transition should be immediately apparent to mice due to the lack of a delay period in each trial. Indeed, performance on control trials in the simple task was as high in mice with delay task experience as in mice with simple task experience only, indicating that subjective task difficulty did not differ depending on training history (Figure 2F). Also, inhibition of S1 had little effect on performance in the simple task both with and without previous delay task training. Therefore, mice with distinct previous task experience require different cortical areas to perform the same task.

**Figure 2.**
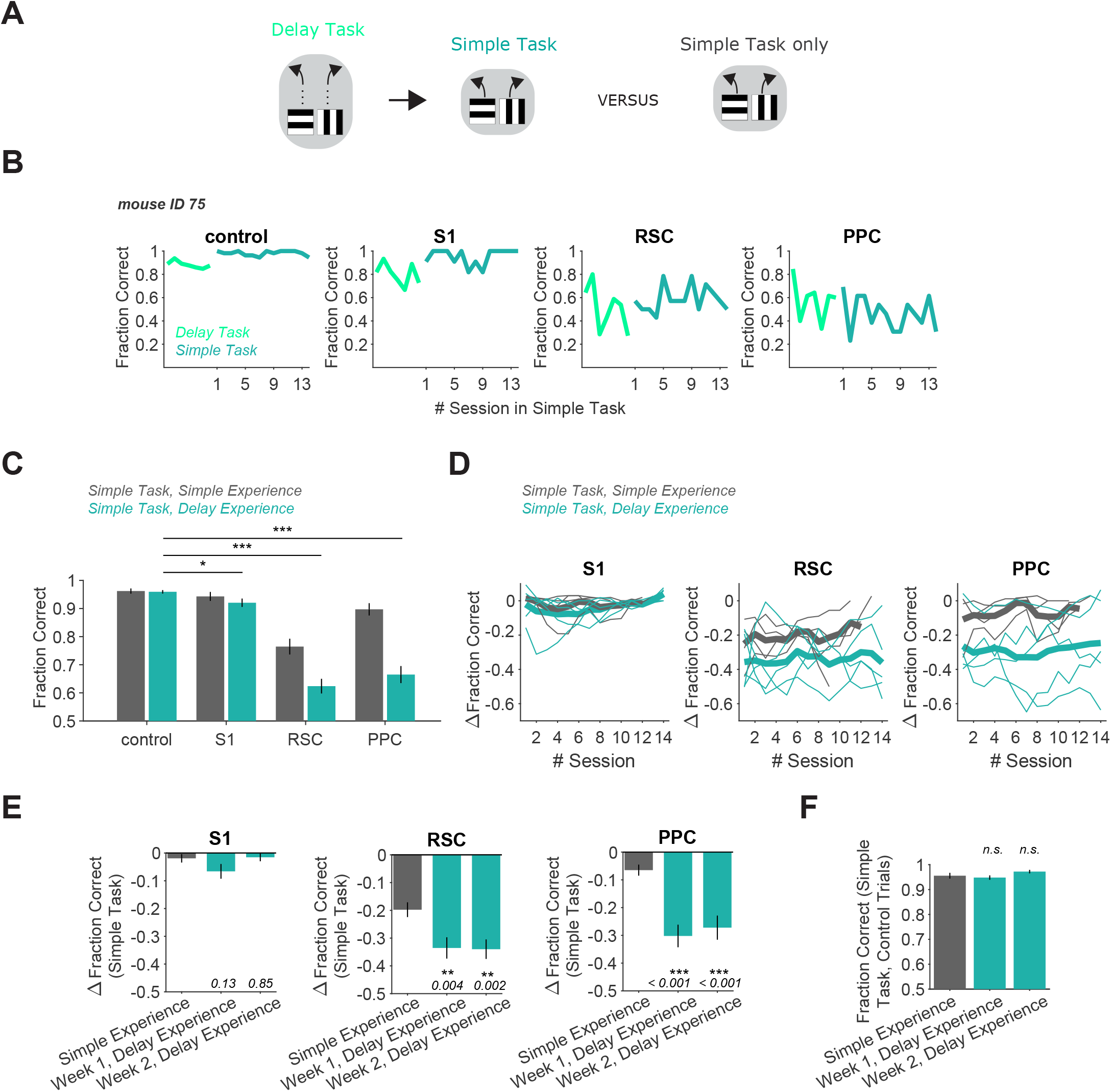
Delay task experience increases the necessity of RSC and PPC in a simple decision task. **(A)** Schematic of the training history sequence. One group of mice was trained on the delay task and then permanently transitioned to the simple task. This group of mice was compared to another group trained only on the simple task. **(B)** Performance of an example mouse transitioned from the delay task to the simple task on control and inhibition trials. **(C)** Performance in the simple task for each inhibited location in mice with simple task experience only (gray, 45 sessions from 4 mice, same dataset as in Figure 1F), and in mice with previous delay task experience (blue, 70 sessions from 5 mice). Bars indicate mean ± sem of a bootstrap distribution of the mean. S1 p = 0.012; RSC p < 0.001; PPC p < 0.001; from bootstrapped distributions of ΔFraction Correct (difference from control performance) compared to 0, two-tailed test, α = 0.05 plus Bonferroni correction. Sessions per mouse: 14. Trials per session: 53 ± 7 (control), 13 ± 3 (S1), 13 ± 2 (RSC), 14 ± 2 (PPC), mean ± SD. **(D)** Inhibition effects (ΔFraction Correct) across sessions in the simple task in mice with only simple task experience (gray), and in mice with previous delay task experience (blue), for each cortical inhibition location. Thin lines show individual mice (n = 4 with simple task experience, n = 5 with delay task experience), thick lines show average across mice. ΔFraction Correct was smoothed with a moving average filter of 3 sessions. **(E)** Comparison of inhibition effects (ΔFraction Correct) in the simple task for mice with simple task experience only (45 sessions from 4 mice) versus delay task experience 1 or 2 weeks after transition from the delay task to the simple task (35 sessions per week from 5 mice). Bars indicate mean ± sem of a bootstrap distribution of the mean; two-tailed comparisons of bootstrapped ΔFraction Correct distributions, α = 0.05. Simple experience datasets are the same as in Figures 1F and Figure 2C. **(F)** Comparison of performance on control trials in the simple task with simple versus delay task experience, using only the first two laser-on blocks in each session. Bars indicate mean ± sem of a bootstrap distribution of the mean. Simple task data in week 1 (p = 0.59) and week 2 (p = 0.19) after transition from the delay task were compared to the simple task only experience data; two-tailed comparisons of bootstrapped Fraction Correct distributions, α = 0.05. Trials per session: 51 ± 23 (simple experience), 51 ± 6 (delay experience, week 1), 53 ± 3 (delay experience, week 2), mean ± SD.

We reached a similar conclusion when we compared mice performing the simple task with and without previous training on the switching task (Figure 3A). In mice that had previously been experts on the switching task, PPC and RSC inhibition resulted in performance of 67 ± 3% and 61 ± 3% correct (mean ± SEM), respectively, on the simple task (Figure 3B-C). As for previous training on the delay task, this effect of PPC and RSC inhibition during the simple task was markedly larger than when these areas were inhibited in mice without previous training on the switching task (Figure 3D-E). PPC activity was necessary even two weeks after the transition to the simple task. RSC’s involvement was greatest in the first week after the transition. Therefore, mice with previous experience in the switching task require PPC and RSC activity to perform the simple task, whereas these areas are largely dispensable during performance of the same simple task in mice without this previous training.

**Figure 3.**
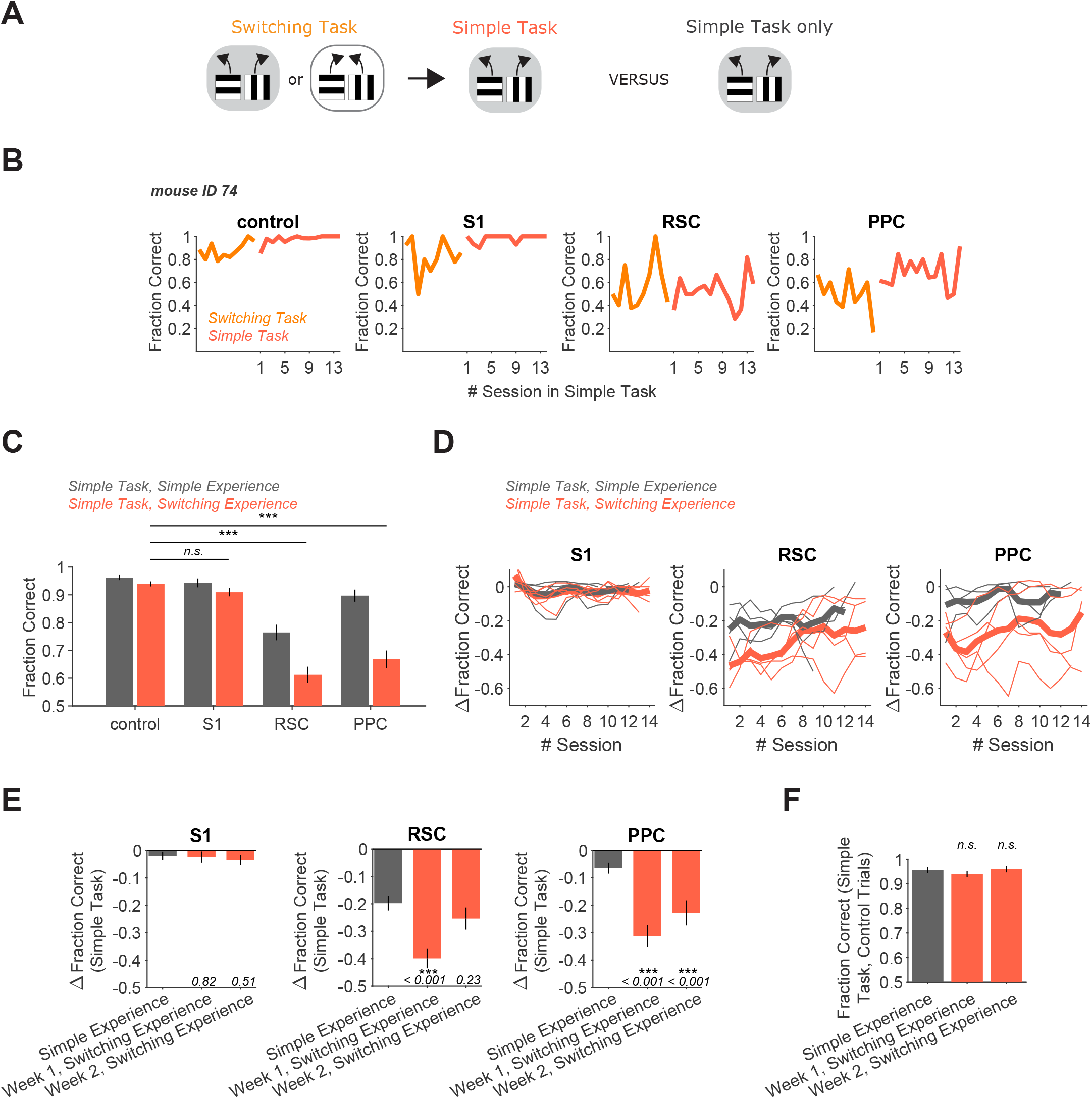
Switching task experience increases the necessity of RSC and PPC in a simple decision task. **(A)** Schematic of the training history sequence. One group of mice was trained on the switching task and then permanently transitioned to the simple task. This group of mice was compared to another group trained only on the simple task. **(B)** Performance of an example mouse transitioned from the switching task to the simple task on control and inhibition trials. **(C)** Performance in the simple task for each inhibited location in mice with simple task experience only (gray, 45 sessions from 4 mice, same dataset as in Figure 1F), and in mice with previous switching task experience (red, 69 sessions from 5 mice). Bars indicate mean ± sem of a bootstrap distribution of the mean. S1 p = 0.26; RSC p < 0.001; PPC p < 0.001; from bootstrapped distributions of ΔFraction Correct (difference from control performance) compared to 0, two-tailed test, α = 0.05 plus Bonferroni correction. Sessions per mouse: 13 ± 0.4. Trials per session: 55 ± 11 (control), 14 ± 4 (S1), 13 ± 4 (RSC), 15 ± 4 (PPC), mean ± SD. **(D)** Inhibition effects (ΔFraction Correct) across sessions in the simple task in mice with only simple task experience (grey), and in mice with previous switching task experience (red), for each cortical inhibition location. Thin lines show individual mice (n = 4 with simple task experience, n = 5 with switching task experience), thick lines show average across mice. ΔFraction Correct was smoothed with a moving average filter of 3 sessions. **(E)** Comparison of inhibition effects (ΔFraction Correct) in the simple task for mice with simple task experience only (45 sessions from 4 mice) versus switching task experience 1 (35 sessions from 5 mice) or 2 (34 sessions from 5 mice) weeks after transition from the switching task to the simple task. Bars indicate mean ± sem of a bootstrap distribution of the mean; two-tailed comparisons of bootstrapped ΔFraction Correct distributions, α = 0.05. Same datasets as in Figures 1F and Figure 3C. **(F)** Comparison of performance on control trials in the simple task with simple versus switching task experience using only the first two laser-on blocks in each session. Bars indicate mean ± sem of a bootstrap distribution of the mean. Simple task data in week 1 (p = 0.32) and week 2 (p = 0.81) after transition from the switching task were compared to the simple task only experience data; two-tailed comparisons of bootstrapped Fraction Correct distributions, α = 0.05. Trials per session: 51 ± 23 (simple experience), 51 ± 5 (switching experience, week 1), 50 ± 7 (switching experience, week 2), mean ± SD.

We wondered if the mice transitioned from the switching task to the simple task might continue to behave as if they were in the dynamic context of the switching task. However, it appeared that mice adapted behaviorally to the simple task quickly after the transition. First, performance on control trials improved to levels observed in mice without previous training on the switching task (Figure 3F). Also, in the switching task, performance at the start of sessions was only at intermediate levels as mice determined the current rule (Figure 3—figure supplement 1). In contrast, after a few days in the simple task, performance was near perfect even in the first tens of trials within a session. Interestingly, when presented with the opposite rule from the switching task again after two weeks on the simple task, mice could still switch back to the long unseen rule within a single session (Figure 3—figure supplement 1). Thus, although mice appeared to retain an understanding of potential association switches, their behavior did not reflect such expectations soon after they were transitioned to the simple task.

We also assessed whether the persistent increase of cortical necessity due to complex-task experience extended to the run-to-target task. Notably, inhibition of PPC and RSC during the run-to-target task resulted in similarly minor performance drops in groups of mice with and without prior complex-task experience (Figure 3—figure supplement 2). Thus, previous experience did not make cortex essential for all tasks.

Collectively, these results highlight that the cortical areas used to perform a task can be profoundly shaped by experience from weeks ago. Mice with different previous task experience use distinct sets of cortical areas to solve the same task. Therefore, an understanding of which areas of cortex are necessary for decision tasks requires considering both the demands of the task-of-interest and the previous experiences of the individual.

### PPC and RSC neurons have activity patterns with higher selectivity in the switching task

Given that the necessity of cortical areas was modulated by previous training, we next asked if the neural activity patterns in these areas are also affected. One possibility is that the increased necessity of cortical areas as a result of previous learning is due to changes in the neural activity patterns within the area. Alternatively, previous learning might not affect a cortical area’s activity, and instead the change in a cortical area’s necessity could be due solely to how its activity is read out by downstream areas (Chowdhury & DeAngelis, 2008; Liu & Pack, 2017).

We simultaneously measured the activity of neurons in PPC, RSC, and V1 with two-photon calcium imaging using a large field-of-view, random-access microscope (Sofroniew et al., 2016) (Figure 4A-C). This microscope allowed us to simultaneously image hundreds of neurons in each of these three cortical areas, with single-cell resolution. We focused our imaging on PPC and RSC because these areas showed major differences in necessity for decisions depending on previous experience. We also included V1 because it is densely interconnected with PPC and RSC (Zhang et al., 2016) and is likely necessary for visual navigation tasks, at least for visual processing (Resulaj et al., 2018). We restricted our imaging experiments to the simple task and the switching task, given that they contain the identical virtual environments.

**Figure 4.**
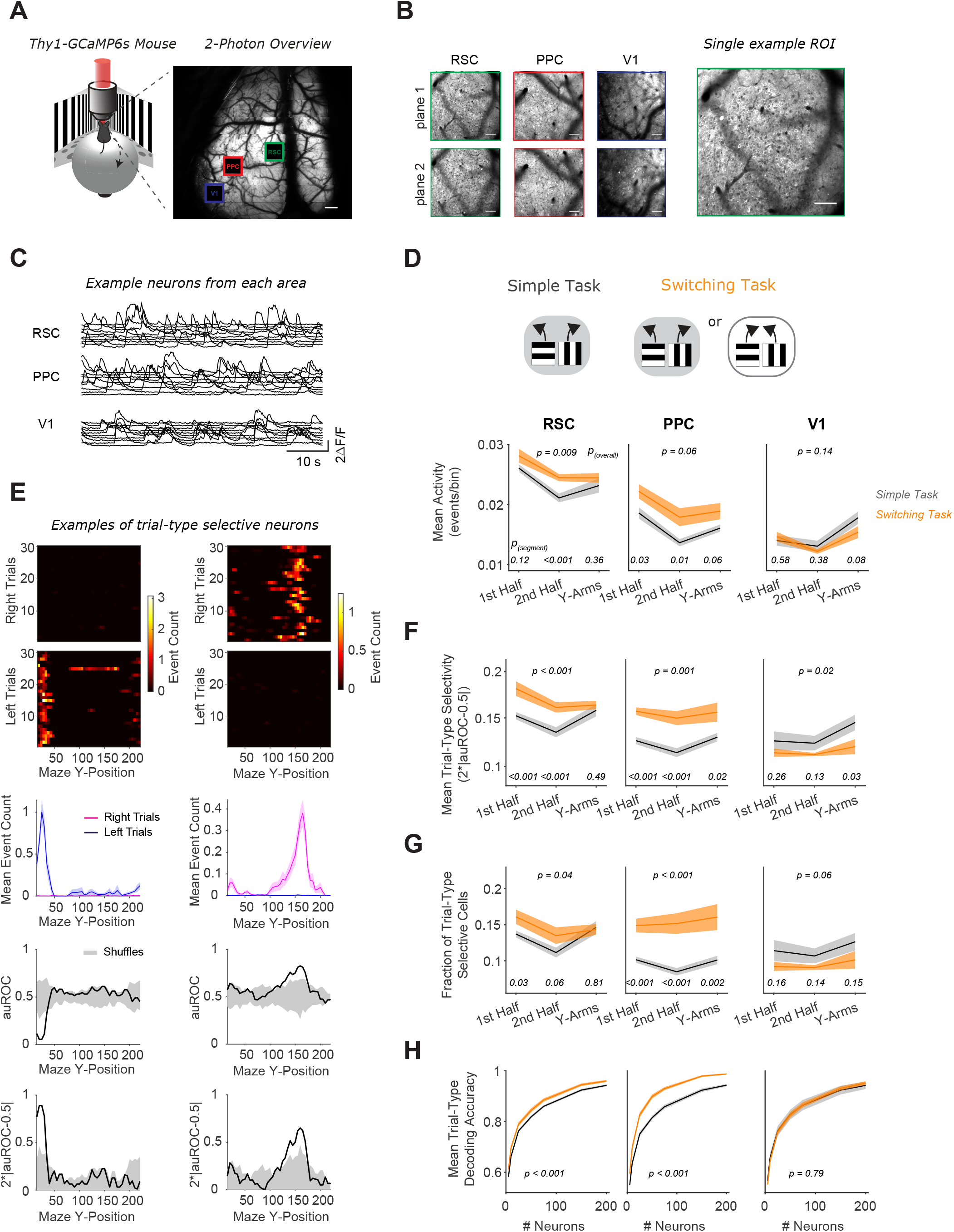
RSC and PPC neurons have activity patterns with higher selectivity in the switching task. **(A)** Left: Schematic of virtual reality behavioral setup with mesoscopic two-photon imaging. Right: Two-photon overview image of cortical window and locations of three areas imaged simultaneously. Scale bar: 500 μm. **(B)** Mean intensity two-photon images of each imaged area, color-coded by area as in **(A)**. Areas were imaged at two depths. Scale bars: 100 μm. **(C)** Example activity traces of ten cells from each area. **(D)** In each area, mean activity levels across cells by maze segment are compared in the simple (gray) versus the switching (orange) task (only Rule A trials). The maze segments are the first half of the Y-stem (15-75 cm Y-Position), the second half of the Y-stem (75-150 cm), and the Y-arms (150-220 cm). Shading indicates mean ± sem of bootstrapped distributions of the mean. P_(segment)_ shows p values of two-tailed comparisons of bootstrapped distributions per maze segment. P_(overall)_ shows the p value for the task factor from a two-way ANOVA (factors: task and maze segment). Simple task: n = 3 mice, 4 sessions per mouse, neurons per session by area: RSC: 1438 ± 217, PPC: 456 ± 172, V1: 498 ± 170. Switching task: n = 3 mice, 4 sessions per mouse, neurons per session by area: RSC: 1510 ± 398, PPC: 513 ± 304, V1: 363 ± 92 (mean ± SD). **(E)** Left and right panel columns show activity from two different example neurons. Top: Spatially binned activity separated by trial type. Each row shows a single trial. Trials were subsampled to 30 trials per trial type. Middle: Mean activity for each trial type (pink: right trials, blue: left trials). Bottom: The area under the ROC curve (auROC) was calculated for each spatial bin. A shuffled distribution of auROC values (gray) was generated by randomly assigning left/right trial labels to each trial and recomputing auROC 100 times. Trial-type selectivity was defined as an absolute deviation of auROC from chance level (2*|auROC-0.5|). To determine significance of trial-type selectivity, at each bin, this value was compared to the trial-type selectivity of the shuffle distribution (gray, significance threshold of p < 0.01). **(F)** Similar to **(D)**, except for the metric of trial-type selectivity, i.e. 2*|auROC-0.5|. **(G)** Similar to **(D)**, except for the fraction of trial-type selective cells as determined from comparing each cell’s selectivity value to a distribution with shuffled trial labels (significance threshold of p < 0.01). **(H)** In each area, trial-type decoding accuracy using activity of subsampled neurons is compared in the simple versus the switching task (Rule A trials only). Shading indicates mean ± sem across sessions. p value is for the task factor from a two-way ANOVA (factors: task and neuron number).

We first imaged neural activity in separate sets of mice in the simple task and switching task (Figure 4D). To compare identical trial types across tasks, that is trials with the same cue-choice associations, we compared neural activity in mice performing the simple task to activity specifically in Rule A of mice performing the switching task. In the switching task, we only included trials once performance had recovered to high levels after rule switches. We started by looking at a basic measure of neural activity, the overall level of activity in individual neurons. Interestingly, this basic measure revealed differences across tasks, as neurons in RSC and PPC had higher activity in the switching task than in the simple task (Figure 4D).

Next, we considered that a direct way a cortical area may contribute to a decision task is by having activity that is different for the two trial types containing distinct cue-choice associations. Trial-type selectivity is a common measure for neural correlates of decision-related functions because it would allow a downstream area to read out the identity of the association and to execute the appropriate choice. We measured this selectivity as our ability to identify the trial type based on a neuron’s activity and quantified it as the area under the receiver operating characteristics curve (auROC, Figure 4E). PPC and RSC neurons showed higher average levels of trial-type selectivity in the switching task than in the simple task (Figure 4F). At the level of populations of neurons, the trial-type selectivity was structured as sequences of neural activity, in which individual neurons were transiently active and different neurons were active at different locations along the maze (Figure 4—figure supplement 1). In addition, the fraction of RSC or PPC neurons with significant trial-type selectivity was higher in the switching task than in the simple task (Figure 4G).

We next assessed how well the current trial type could be decoded from the activity of neural populations of varying sizes in each area. In the simple task, RSC and PPC populations contained task-relevant information that led to above-chance decoding from a population of neurons. However, for the same size population in the switching task, this decoding accuracy in RSC and PPC was even higher, in line with the observed increased selectivity and larger fraction of selective neurons relative to the simple task (Figure 4H). These differences in activity across tasks in RSC and PPC were especially striking because, in the simple task and Rule A of the switching task, mice ran through a maze with identical visual cues and made similar left-right behavioral choices in both tasks. Therefore, the activity levels and selectivity of single neurons are higher in PPC and RSC when mice perform a more complex task, even when the sensory stimuli and choice reports in the tasks are identical, leading to better ability to decode the trial type.

We verified that the differences in selectivity across tasks were not due to differences in running patterns. When we selected sessions so that the time course and magnitude of decoding the mouse’s reported choice from its running were similar across tasks, we largely observed the same differences in neural trial-type selectivity as reported above (Figure 4—figure supplement 2). Thus, the differences in neural selectivity cannot be trivially explained by differences in running patterns.

In contrast, V1 neurons had similar levels of activity, selectivity, and population-level trial-type decoding in the simple and switching tasks (Figure 4D-H). This finding is consistent with the identical visual scene in these tasks but is perhaps surprising given that V1 neurons have been shown to contain many non-visual signals (Koay et al., 2020).

### Previous switching task experience increases neural trial-type selectivity in PPC and RSC

We then examined if previous experience in the switching task affected the neural activity patterns in the simple task. Similar to our tests of cortical necessity, we compared neural activity during the simple task in mice that either had or had not been trained previously in the switching task (Figure 5A). We trained one group of mice on the switching task and then permanently transitioned these mice to the simple task. A separate set of mice was trained only on the simple task. We thus compared the activity patterns in PPC, RSC, and V1 in mice performing decisions in the same task except with distinct experience.

**Figure 5.**
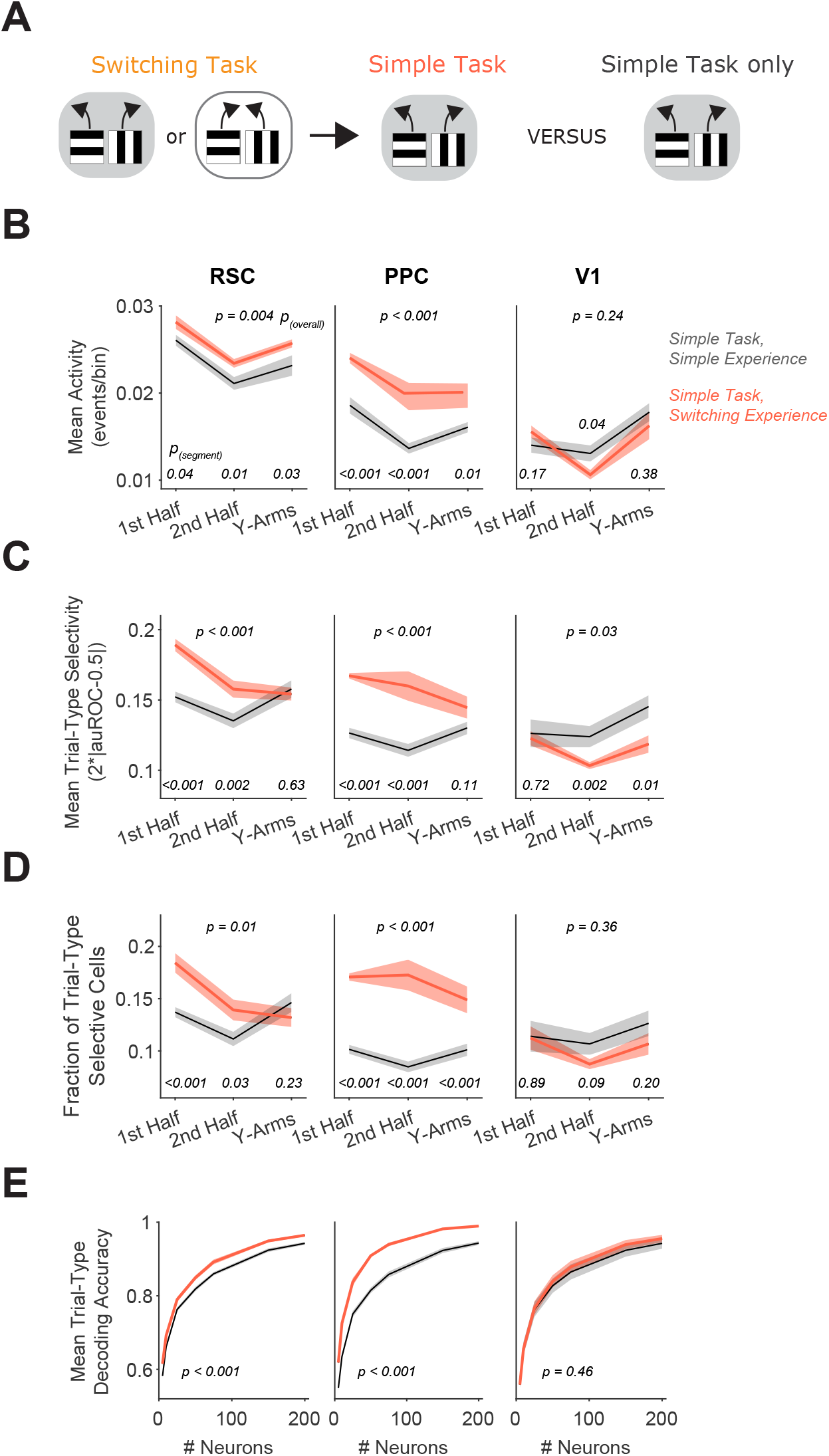
Previous switching task experience increases trial-type selectivity in RSC and PPC. **(A)** Schematic of the training history sequence. One group of mice was trained on the switching task and then permanently transitioned to the simple task. This group of mice was compared to another group trained only on the simple task. **(B)** In each area, mean activity levels across cells by maze segment are compared in the simple task in mice with (red) and without (gray) previous experience in the switching task. Shading indicates mean ± sem of bootstrapped distributions of the mean. P_(segment)_ shows p values of two-tailed comparisons of bootstrapped distributions per maze segment. P_(overall)_ shows the p value for the previous task experience factor from a two-way ANOVA (factors: previous task experience and maze segment). Simple task: n = 3 mice, 4 sessions per mouse, cells per session by area: RSC: 1438 ± 217, PPC: 456 ± 172, V1: 498 ± 170 (same dataset as in Figure 4D-H). Simple task after switching task experience: n = 2 mice, 3 and 5 sessions per mouse, neurons per session by area: RSC: 1407 ± 327, PPC: 744 ± 219, V1: 351 ± 90 (mean ± SD). **(C)** Similar to **(B)**, except for the metric of trial-type selectivity, i.e. 2*|auROC-0.5|. **(D)** Similar to **(B)**, except for the fraction of trial-type selective cells as determined from comparing each cell’s selectivity value to a distribution with shuffled trial labels. **(E)** In each area, trial-type decoding accuracy using activity of subsampled neurons is compared in the simple task in mice with and without previous experience in the switching task. Shading indicates mean ± sem across sessions. p value is for the previous task experience factor from a two-way ANOVA (factors: previous task experience and neuron number).

Strikingly, during the simple task, neurons in RSC and PPC had higher activity in mice with experience in the switching task than in mice trained only in the simple task (Figure 5B). Furthermore, in mice with switching task experience, RSC and PPC neurons had higher average selectivity for the trial type and higher fractions of neurons with significant trial-type selectivity (Figure 5C-D). As a result, the decoding of the trial type from population activity was more accurate in these areas in mice with the complex task experience (Figure 5E). Notably, selectivity in V1 neurons was similar between mice with and without complex task experience. Therefore, the activity patterns of single neurons in PPC and RSC, including mean activity levels and selectivity, are strongly influenced by previous task experience.

### Switching task experience decreases noise correlations

A key feature of neural codes beyond the properties of single cells is the collective activity of populations of neurons. Properties of population codes affect the amount of information in neural populations and have been shown to depend on behavioral context, task learning, and other factors (Cohen & Kohn, 2011). We thus examined if features of the neural population code are also affected by task complexity and past experience. We took advantage of the simultaneous recording of hundreds of neurons per area and analyzed the correlation in activity for pairs of neurons on a given trial type, a measure commonly referred to as noise correlation, that quantifies trial-to-trial co-fluctuations in neurons (Cohen & Kohn, 2011). As for analyses of trial-type selectivity, we restricted analyses to the maze traversal period and only included correct trials from high performance periods (see Methods). The noise correlations within individual areas and across pairs of areas were lower on average in mice performing the switching task than in mice performing the simple task (Figure 6A-C). Strikingly, when we compared mice with different experience as they performed the same simple task, we also observed a difference in noise correlations both within and across cortical areas, with lower correlations in mice with switching task experience compared to mice trained only in the simple task (Figure 6A-C). Therefore, mice with different task experience have significant differences in their population codes as they perform the same task.

**Figure 6.**
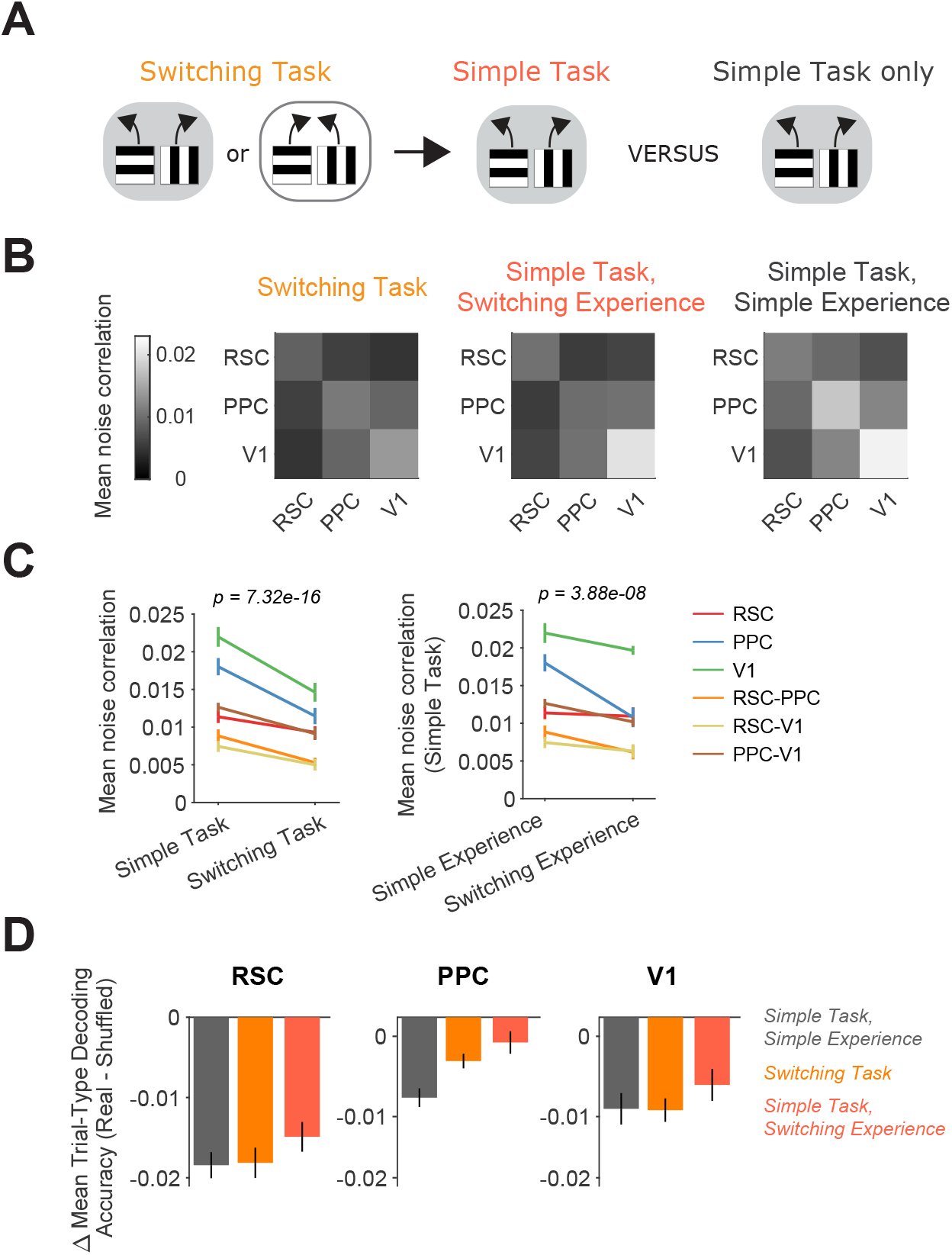
Switching task experience decreases noise correlations. **(A)** Schematic of the training history sequence. One group of mice was first trained on the switching task and then permanently transitioned to the simple task. Another group was trained only on the simple task. **(B)** Mean pairwise noise correlations within and across areas from bootstrapped distributions of the mean in the switching task (left), the simple task with previous switching task experience (middle), and the simple task with only simple task experience (right). Noise correlations were calculated on spatially binned data in correct trials during high performance periods (Methods). **(C)** Left: Comparison of mean noise correlations in the switching task versus the simple task, p value is for the task factor from a two-way ANOVA (factors: task and area-combination). Error bars indicate mean ± sem across sessions per area combination (n = 6 area combinations, n sessions: 12 (simple task), 12 (switching task)). Right: Similar to left, except for the comparison of simple task noise correlations with and without previous switching task experience. n sessions: 12 (simple experience), 8 (switching experience). **(D)** For each area and task or previous task experience, the difference in trial-type decoding accuracy between neural populations (200 subsampled neurons) with intact and disrupted noise correlations. Noise correlations were disrupted by shuffling trials independently for each cell within a given trial type. Error bars show mean ± sem across sessions.

Noise correlations can in some cases have detrimental effects on population codes by limiting the information capacity because these correlations are co-fluctuations in activity that cannot be removed by averaging across neurons (Averbeck et al., 2006; Kafashan et al., 2021; Panzeri et al., 1999; Zohary et al., 1994). To reveal the impact of correlations on coding in our experiments, we disrupted noise correlations by shuffling trials of a given trial type separately for each neuron and repeated the trial-type decoding. The accuracy of decoding the trial type was slightly higher with correlations disrupted (Figure 6D). Therefore, the lower correlations in mice performing the switching task or the simple task with switching task experience boosts information encoding along with higher trial-type selectivity levels.

Thus, these results reveal that the activity patterns in single neurons and neural populations are shaped by previous experience. Together, our findings demonstrate that different sets of cortical areas and distinct neural activity patterns are utilized for the same task depending on an individual’s training history.

## Discussion

We have shown that the same task, with the same visual cues and behavioral choice reports, is solved using distinct cortical areas and activity patterns depending on the past experience of the individual. Thus, the necessity of cortical areas for a decision task depends on factors separate from the task itself. Here, we aimed to vary the “cognitive experience” of the mouse, which we define as the features of the task other than the choice-informative sensory stimuli and the behavioral outputs. We varied the cognitive experience by adding delay periods or frequent switches of associations within a session, but the maze shape, and thus behavioral outputs needed, and choice-informative cues were identical between the complex and simple tasks. Thus, the differences between mice with and without training on the complex tasks is likely due to cognitive experience instead of sensory or motor learning. Our results reveal that mice with enhanced cognitive experience due to previous training on complex tasks require PPC and RSC to perform simple decision tasks. In contrast, in mice without this previous training, PPC and RSC are largely dispensable for performing the same, simple task. This difference in cortical necessity was accompanied by differences in the activity patterns of single neurons and neural populations. During the same task, mice with complex training experience had higher selectivity in single neurons and weakened neuron-neuron correlations, compared to mice without this history, which together allowed for easier decoding of the relevant information from a population of neurons. Together, these results show that distinct sets of cortical areas can be used to solve the same task.

We specifically varied the cognitive experience of mice while keeping the sensory cues and behavioral choice reports the same across decisions, with a focus on cortical association areas. Previous studies of perceptual experience and motor learning have emphasized that cortical necessity decreases with experience (Chowdhury & DeAngelis, 2008; Hwang et al., 2019; Kawai et al., 2015) (but see (Liu & Pack, 2017)). Instead, we found an increased necessity of cortical association areas for simple decisions due to cognitive experience in two distinct, complex tasks (delay and switching tasks). Future work should aim to understand how differences in the type of experience (cognitive versus perceptual), types of tasks, and areas studied (association versus sensory or motor cortices) influence whether experience increases or decreases cortical necessity.

In addition, these earlier works did not identify differences in neural tuning in MT with perceptual experience despite differences in necessity, leading to the proposal that differences in cortical necessity with experience arise from whether an area’s activity is read out by a downstream network. Here, we did find differences in neural selectivity specifically in association areas. However, these areas already contained task-relevant information in the simple task without complex task experience. It appears unlikely that the observed boost in neural selectivity with complex task experience is the sole reason for the large increase in cortical necessity for task performance. Increased cortical necessity may rather result from a combination of increases in task information, reshaped representations of information, and/or modifications to information readouts (Ruff & Cohen, 2019). Further work will be needed to test directly the relationship between specific features of the neural code and the causal roles of PPC and RSC in decision tasks.

We found that cortical association areas had higher selectivity in their neural activity and were more strongly required for the complex tasks than the simple task. This finding supports the notion that cortex is needed for cognitively more challenging tasks. Previous work presented a similar finding but focused on tasks with a wide gap in their demands, such as comparing running toward a visual target versus using learned cue-choice associations to make navigation decisions (Buschman et al., 2011; Ceballo et al., 2019; Fuster, 1997; Harvey et al., 2012; Lashley, 1931; Pinto et al., 2019; Sarma et al., 2015). In our work, we extended this concept by keeping the visual and behavioral aspects of the complex and simple tasks as similar as possible and adding specific cognitive challenges. Our goal here was not to compare the specific features of the different tasks, and instead we used the complex tasks to establish different previous experiences. Future work will be needed to compare in more depth the neural activity in the different tasks and to understand how PPC and RSC might contribute to the switching and delay tasks.

Our results have crucial implications for the experimental study of cortical involvement in decision-making. Many studies, including our previous work, test an area’s involvement or activity patterns in a single task and develop interpretations of an area’s functions by extrapolating across studies (Lyamzin & Benucci, 2019). However, given that factors beyond the task-of-interest contribute to an area’s necessity and activity in a task, we encourage consideration of a variety of factors that have often been ignored and not reported. Because prior task expertise can have a large effect on the involvement of cortical areas, it seems critical to report the full details of how animals were trained on a task and what previous experiences they encountered, both of which are commonly omitted from publications.

Together, our results indicate that the cortical implementation of a decision task flexibly depends on initial conditions, as defined by past experience, and the overall optimization goal for the animal, which in many cases is not just a single task but also previous tasks or other tasks occurring in parallel (Golub et al., 2018; Sadtler et al., 2014). These results highlight the tremendous flexibility of the brain to perform outwardly identical tasks using distinct sets of brain areas and neural activity patterns and raise exciting challenges for understanding neural computation in the framework of dynamic and distinct neural solutions for a given cognitive problem. We propose that understanding cognitive processes will require considering the wider set of functions an animal is trying to optimize, beyond the decision or computation of interest in a particular study. To understand how long-ago cognitive experience and current cognitive demands set up these different neural circuit landscapes for outwardly identical decisions, we suggest to carefully control for and to intentionally vary cognitive experience in laboratory settings (Plitt & Giocomo, 2021; Sharpe et al., 2021), thereby approximating more naturalistic decision-making scenarios. We anticipate this approach will be particularly powerful to illuminate neural circuit differences underlying inter-individual variability and changes in neural dynamics during learning (Oby et al., 2019; Sadtler et al., 2014).

## Acknowledgements

We thank Yvette Fisher, Lauren Orefice, and members of the Harvey lab for feedback on the manuscript, Matthias Minderer for optogenetics designs and code, Shih-Yi Tseng for input on behavioral training, Kıvılcım Kılıç for demonstrating a crystal skull cranial window surgery, and the Research Instrumentation Core at Harvard Medical School. This work was supported by grants from the NIH (R01 MH107620, R01 NS089521, R01 NS108410, DP1 MH125776), an NIMH Diversity Supplement, a Louis Perry Jones Postdoctoral Fellowship (CA), Alice and Joseph Brooks Postdoctoral Fellowships (CA, SK), a Mahoney Postdoctoral Fellowship (CA), a Leonard and Isabelle Goldenson Postdoctoral Fellowship (SK), a Uehara Foundation Research Fellowship (SK), a NARSAD Young Investigator Grant (SK), a JSPS Overseas Research Fellowship (SK), an EMBO postdoctoral fellowship (SS), and a Stuart H.Q. & Victoria Quan Fellowship (NLP).

## Author Contributions

CA and CDH conceived of the project. CA, RBL, and CDH designed the experiments. CA and RBL performed the experiments with assistance from SK, CAB, and NX. CA analyzed the data with assistance from RBL and input from CDH. SNC developed the switching task. NLP developed the compact virtual reality design. CA and SS built the random-access two-photon microscope. CDH oversaw all aspects of the research and obtained funding for the research. CA and CDH wrote the paper with input from all authors.

## Data and Materials Availability

Data and custom code will be made available upon publication.

## METHODS

### Mice

All experimental procedures were approved by the Harvard Medical School Institutional Animal Care and Use Committee and were performed in compliance with the Guide for the Care and Use of Laboratory Animals. All optogenetic inhibition data were acquired from 19 male VGAT-ChR2-YFP mice (The Jackson Laboratory, stock 014548). All calcium imaging data were acquired from six C57BL/6J-Tg(Thy1-GCaMP6s) GP4.3Dkim/J mice (stock 024275) of both sexes (5 female, 1 male). Mice were 11-32 weeks old at the start of behavioral training. Age at training start did not vary systematically across tasks (simple task: 21 ± 7 weeks (n = 7), switching task: 17 ± 5 weeks (n = 13), delay task: 21 ± 11 weeks (n = 5), mean ± SD, including photoinhibition and calcium imaging mice). Mice were kept on a reverse dark/light cycle and housed in groups of 2-3 littermates per cage (mice for optogenetic inhibition) or single-housed (mice for calcium imaging).

### Virtual reality setup

Virtual reality environments (Harvey et al., 2009) were designed and operated in VirRMEn (Virtual Reality Mouse Engine) (Aronov & Tank, 2014). A novel, custom-made compact virtual reality design was employed (overall dimensions of approximately 15 inches wide x 21 deep x 18 high), using modifiable laser-cut acrylic and mirror pieces. A micro projector (Laser Beam Pro) projected the virtual environment onto a double-mirror system and a 15-inch diameter half-cylindrical screen. The mouse was head-fixed on top of an 8-inch diameter Styrofoam spherical treadmill. The mouse’s position in the virtual environment, and thus the projection, was controlled by the mouse’s movement of the treadmill, which was measured with two optical sensors (ADNS-9800, Avago Technologies) placed 90**°** apart from each other beneath the ball. The treadmill velocity was translated into pitch, roll, and yaw velocity relative to the mouse’s body axis using custom code on a Teensy microcontroller (PJRC). Pitch controlled forward/backward movement in the virtual world, while roll controlled lateral movement. The virtual view angle was fixed so that the mouse could not rotate in the virtual world. Designs for the virtual reality apparatus are available at https://github.com/HarveyLab/mouseVR.

### Behavioral training

Mice were limited to 1 mL of water per day for several days before starting training. Body weight and body condition were checked daily. Mice were maintained at approximately 80% of their body weight prior to water restriction and received additional water if their body weight fell below 75% of their original weight. Mice were trained daily for 45-80 minutes (except for some weekends, when they received 1 mL of water without training). In the first training phase, to get accustomed to the experimental setup and to moving in virtual space, mice had to run down the length of a rectangular environment (‘linear maze’) towards a checkerboard pattern. When reaching the checkerboard, they received a reward (3-4 μL of 10% sweetened condensed milk in water delivered through a lick spout) and were teleported back to the beginning after a brief inter-trial-interval (ITI, 1 s). The linear maze contained the visual cues on the walls that the mice would later learn to associate with rewarded choice directions. Linear maze length was increased on a session-by-session basis until mice completed approximately 200 trials of a 200 cm maze in 60 minutes (minimum / maximum maze length of 10 cm / 200 cm).

#### Y-Maze training, general procedures

After training on the linear maze, mice were transitioned to a Y-shaped maze (180 cm long) in which they had to run towards one of two possible Y-arms to get rewarded. In all tasks, visual cues presented on maze walls were associated with rewarded choice arms at the end of the maze. Initially, in all tasks, the correct choice was signaled with a checkerboard at the end of the correct Y-arm (100% “visually guided trials”). Throughout training, the fraction of “visually guided trials” was gradually reduced on a session-by-session basis by the experimenter, based on the mouse’s previous performance. For the simple and delay task, average performance in the preceding session had to exceed 80%. For the switching task, overall performance including periods after rule switches had to exceed 70% correct, performance after rule switches was inspected for drops followed by recovery over tens of trials, and mice had to obtain at least two rule switches per session. Across all tasks, after extensive training, a minority of trials were still “visually guided trials” (10-15%). In non-visually guided trials, the checkerboard appeared for 2 s as visual feedback once the mouse made a correct choice, followed by the reward and an ITI with a gray screen of 2 s. After incorrect choices, no checkerboard was presented, and the ITI lasted for 4 s. A different cohort of mice was trained on each task unless specified otherwise. As some mice developed biases during training, making predominantly left or right choices, we employed a bias correction algorithm in some training sessions. If the mouse made the same choice on the last five trials irrespective of correctness, the next trial would contain a maze in which the opposite choice would be correct. This bias correction was only used at intermediate training stages and was not employed during inhibition or calcium imaging sessions.

#### Simple task

In the simple task, mice encountered one of two possible cues in a given trial, with a fixed association between cue identity and rewarded Y-arm. Horizontal gratings were associated with a left rewarded choice and vertical gratings were associated with a right rewarded choice. This pair of associations constitutes what we call ‘Rule A’ in the switching task. Visual cues were present along the entire extent of the maze, including the stem and the Y-arms.

#### Delay task

Mice trained in the delay task were first trained in the simple task. After reaching high performance levels in the simple task (at least 90% correct), a delay was introduced, i.e. a neutral visual texture present in all trial types that was uninformative about the choice to make on a given trial. In the first sessions of delay task training, the delay texture was only present in the Y-arms of the maze. Then the delay onset (i.e. cue offset) was gradually shifted earlier in the trial by 10 cm increments on a session-by-session basis if performance in the preceding session exceeded 80% correct, until only the first half of the Y-maze stem contained the visual cue (50 cm). Thus, mice had to traverse the rest of the maze without the informative cue on the walls. A subset of mice (2 out of 7: Mouse IDs 38, 41) were used for photoinhibition experiments both during the simple task before delay task training, and later during the delay task after delay task training.

#### Switching task

In the switching task, mice encountered the trial types that were visually identical to those in the simple task, but now the associations between visual cue and rewarded choice were switched in blocks (Rule A and Rule B). In ‘Rule B’, mice had to make the opposite choices given the same cue identities as in Rule A to get rewarded, i.e. horizontal / vertical gratings were associated with a rightward / leftward choice, respectively. Rule switches were not explicitly signaled to the mice, so they had to integrate information of past cues, choices, and rewards to inform their belief about the current rule. Rule switches were present from the first day of the Y-maze training period. Given the increased cognitive demand of the switching task, the fraction of visually guided trials was reduced more slowly in the switching task than in the simple task. The initial rule in a given session was alternated on a daily basis, starting out with Rule A on the first day of training. Within a training session, a rule switch occurred if several criteria were met: a minimum of 75 trials from the previous rule switch or the session start, a minimum average performance in the last 30 trials of 85% correct, and a correct choice on the immediately preceding trial. Mice encountered 2-3 rule switches per session, indicating they could repeatedly switch associations successfully.

#### Run-to-visual-target task

To establish a baseline for cortical involvement in a simple navigation task in which mice did not use visual cues on the maze walls to guide their choices, we employed a run-to-visual target task. The maze had the standard Y-shape architecture, but no informative visual cues on the maze walls. Instead, mice simply had to run towards the checkerboard present at one of the two Y-arm ends in each trial. The checkerboard location (left or right) was randomly chosen on each trial. We used mice previously trained in either the simple task only (Figure 1—figure supplement 1) or trained on a complex task (switching or delay) before simple-task-only exposure (Figure 3—figure supplement 2), so the mice already knew that the checkerboard signified a reward location and required minimal training on this task (1-2 days prior to photoinhibition).

### Photoinhibition experiments

#### Clear skull cap surgery

We followed procedures described previously (Guo et al., 2014; Minderer et al., 2019). In brief, the scalp and the periosteum were removed from the dorsal skull surface. The skull surface was covered with a thin layer of cyanoacrylate glue (Insta-Cure, Bob Smith Industries). A bar-shaped titanium headplate was attached to the interparietal bone using dental cement (Metabond, Parkell). Several layers of transparent dental acrylic (Jet Repair Acrylic, Lang Dental, P/N 1223-clear) were applied to the parietal and frontal bones to create a transparent skull cap. In a subsequent procedure preceding photoinhibition experiments, the acrylic was polished with a polishing drill (Model 6100, Vogue Professional) with denture polishing bits (HP0412, AZDENT). Clear nail polish was applied on top of the polished acrylic (Electron Microscopy Sciences, 72180). An aluminum ring was attached to the skull using dental cement mixed with carbon powder (Sigma-Aldrich) for light-shielding.

#### Experimental setup and logic

Light from a 470 nm collimated laser (LRD-0470-PFFD-00200, Laserglow Technologies) was focused onto the skull using an achromatic doublet lens (f = 300 mm, AC508-300-A-ML, Thorlabs). We coupled the laser to a pair of galvanometric scan mirrors (6210H, Cambridge Technology) in combination with rapid analog laser power modulation to allow fast movement of the focused beam between cortical target sites. At the focus, the laser beam had a diameter of approximately 200 µm.

We started photoinhibition only after mice reached expert performance in a given task (criterion for the simple or delay task: performance of approximately 85% correct or higher, criteria in the switching task: at least two rule switches with only 15% visually-guided trials per session). We thus started inhibition after shorter training times in the simple task group of mice, compared to the complex task (delay or switching) groups that required longer training times to reach expert performance (see Figure 1). In each session, we bilaterally targeted PPC, RSC, S1, and a control site outside of the brain (on the dental cement) on separate, interleaved trials. For PPC, S1, and control targets, we used single bilateral laser spots, with laser power sinusoidally modulated at 40 Hz and a time-average power of approximately 6.5 mW/spot. For RSC, we used three spots on each hemisphere to match the region’s anatomical extent, with laser power sinusoidally modulated at 20 Hz and a mean power of approximately 5 mW/spot. The target coordinates in mm from bregma were: RSC (-3.5, -2.5, -1.5 anterior-posterior (AP); 0.5 medial-laterial (ML)); PPC (-2 AP, 1.75 ML); S1 (-0.5 AP, 2.5 ML); control (2 AP, 5 ML). Based on previous calibration studies (Guo et al., 2014; Pinto et al., 2019), we estimate that the laser powers employed here inhibited a cortical area with a radius of 1-2 mm per inhibition spot.

In each experimental session, blocks of at least 50 trials without laser light were alternated with laser-on blocks of 50 trials. Laser-on blocks only started if the mouse’s average performance in the preceding 30 trials was at least 85% correct. Thus, in the switching task, laser-on blocks occurred once the mouse had reached stable performance in the current rule block. In the switching task, rule switches happened after the end of each laser-on block. Within laser-on blocks, approximately 50% of trials were control trials, and the laser target location was randomly chosen for each trial. Within a trial, the laser was on from 0.5 s before visual cue onset at the trial beginning until the mouse reached the end of the maze, excluding the visual feedback and reward / ITI periods of the trial. In the run-to-visual-target task and a subset of sessions in the simple task, a single long block of 200 laser-on trials was delivered after the mouse reached high performance levels, again with approximately 50% control trials randomly interleaved with cortical inhibition trials. In the simple task, inhibition effects did not vary between sessions with laser-on blocks of 50 or 200 trials.

For experiments with maze segment-specific inhibition in the delay task (Figure 1—figure supplement 3), the stem of the Y-maze was doubled in length to 200 cm, and the laser-on period per session was restricted to either only the cue period (maze beginning until delay onset) or the delay period (delay onset until the end of the maze, excluding visual feedback and reward / ITI periods). Cue only and delay only inhibition sessions were generally alternated from day to day.

For experiments with ITI inhibition in the switching task (Figure 1—figure supplement 4), PPC was the only cortical inhibition target. PPC was inhibited during the ITI for 50 consecutive trials following either the first or the second rule switch per session. In all other trials, the laser was steered to the control location during the ITI so that rule switches could not be inferred simply from the presence of laser light. Inhibition started upon the mouse reaching one of the two possible Y-arm maze ends and lasted throughout checkerboard feedback presentation and the ITI / reward delivery.

#### Long-term experimental stability and order of experiments across tasks

To ensure stability of experimental conditions across long times, we maintained constant laser power by measuring maximum power daily and cleaning optics if necessary. To ensure stability of inhibition conditions per mouse, we verified the alignment of the laser beam orientation to the mouse’s skull by creating a laser cross pattern to be centered on Bregma and to be aligned with the mouse’s AP-ML skull axes. To aid in the latter, we added marks on the mouse’s dental cement in AP and ML that the laser cross had to intersect. To change alignment horizontally or vertically, we moved an X-Y stage that the laser apparatus was mounted on. We controlled for rotation by slight adjustments to the posts holding the mouse’s headplate. Experiments in different cohorts across tasks were not systematically interleaved but were clustered in time as we iterated through hypotheses throughout the project. We have several indicators that experimental conditions remained stable and that differences across tasks were not the result of experimental drift. First, data for the switching task were collected in several groups of mice spanning the full range of data collection times, yet the average inhibition effects on performance were similar (group 1 (early) (mouse IDs 24, 27, 42) versus group 2 (late) (mouse IDs 72, 73, 74): ΔFraction correct for Rule A: S1: -9 ± 2 % vs -7 ± 2%, RSC: -31 ± 1% vs -38 ± 4%, PPC: -34 ± 3% vs -30 ± 3 %, mean ± sem). Second, data from the simple task with notably smaller inhibition effects were collected in between these two groups and were partly on overlapping days as the first group. Third, in some mice (Mouse IDs 38, 41), we tested the effect of inhibition in the identical mice in the simple task first, as well as after they learned the delay task, observing large differences in cortical inhibition effects in the same mice within weeks (ΔFraction correct in simple versus delay task: S1: -1 ± 2 % vs -18 ± 18%, RSC: -16 ± 6% vs -36 ± 3%, PPC: -5 ± 2% vs -37 ± 8 %, mean ± sem).

### Calcium imaging experiments

#### Large chronic cranial window surgery

We slightly modified procedures described previously (Kim et al., 2016; Kılıç et al., 2021). Mice were injected with Dexamethasone (3 µg per g body weight) 4-8 h prior to surgery and anesthetized with isoflurane (1-2% in air). A cranial window surgery was performed to either fit a ‘crystal skull’ curved window (LabMaker UG) exposing the dorsal surface of both hemispheres (Kim et al., 2016) or the left hemisphere only (Kılıç et al., 2021), or to fit a stack of custom laser-cut quartz glass coverslips (three coverslips with #1 thickness each (Electron Microscopy Sciences), cut to a ‘D’-shape with maximum dimensions of 5.5 mm medial-lateral and 7.7 mm anterior-posterior, and glued together with UV-curable optical adhesive (Norland Optics NOA65)), exposing the left cortical hemisphere. The skull was kept moist using saline throughout the drilling procedure and soaked in saline for one to two minutes before being lifted. The dura was removed before sealing the window using dental cement (Parkell). A custom titanium headplate was affixed to the skull using dental cement mixed with carbon powder (Sigma-Aldrich) to prevent light contamination. A custom aluminum ring was affixed on top of the headplate using dental cement. During imaging, this ring interfaced with a black rubber balloon enclosing the microscope objective for light-shielding.

#### Calcium imaging setup and data acquisition

Data were collected using a large field of view two-photon microscope assembled as described previously (Sofroniew et al., 2016). In brief, the system consisted of a combination of a fast resonant scan mirror and two large galvanometric scan mirrors allowing for large scan angles. Together with a remote focusing unit to rapidly move the focus depth, this setup enabled random access imaging in a field of view of 5 mm diameter with 1 mm depth. The setup was assembled on a vertically mounted breadboard whose XYZ positions and rotation were controlled electronically via a gantry system (Thorlabs). Thus, to position the imaging objective with respect to the mouse, the position and rotation of the entire microscope were adjusted while the position of the mouse remained fixed. The excitation wavelength was 920 nm, and the average power at the sample was 60-70 mW. The microscope was controlled by ScanImage 2016 (Vidrio Technologies). We imaged in three distinct regions in the left cortical hemisphere: V1, PPC, and RSC. These regions were identified based on retinotopic mapping (see below). In each region, we acquired images in layer 2/3 from two planes spaced 50 µm in depth, at 5.36 Hz per plane at a resolution of 512 x 512 pixels (600 µm x 600 µm). Imaging was performed in expert mice in the simple task, switching task, and simple task after switching task experience (criterion for the simple task: performance of approximately 85% correct or higher, criteria in the switching task: at least two rule switches with only 15% visually-guided trials per session). The stem of the Y-maze was extended by 50% (50 cm) compared to the maze architecture in photoinhibition sessions, resulting in a maze length of 230 cm. Each imaging session lasted 45-80 minutes. During imaging, slow drift of the image was occasionally corrected manually by moving the gantry to align the current image with an image from the beginning of the session. For synchronization of imaging and behavior data, both the imaging and the behavior frame clock were recorded on another computer using Wavesurfer (https://wavesurfer.janelia.org/).

#### Retinotopic mapping for selecting calcium imaging locations

We performed retinotopic mapping in mice used for calcium imaging experiments as previously described (Driscoll et al., 2017; Minderer et al., 2019). Mice were lightly anesthetized with isoflurane (0.7 – 1.2% in air). A tandem-lens macroscope was used in combination with a CMOS camera to image GCaMP fluorescence at 60 Hz (455 nm excitation, 469 nm emission). A periodic spherically corrected black and white checkered moving bar (Marshel et al., 2011) was presented in four movement directions on a gamma-corrected 27 inch IPS LCD monitor (MG279Q, Asus). The monitor was centered in front of the mouse’s right eye at an angle of 30 degrees from the mouse’s midline. To produce retinotopic maps, we calculated the temporal Fourier transform at each pixel of the imaging data and extracted the phase at the stimulus frequency (Kalatsky & Stryker, 2003). These phase images were smoothed with a Gaussian filter (25 μm s.d.). Field sign maps were generated by computing the sine of the angle between the gradients of the average horizontal and vertical retinotopic maps.

For each retinotopic mapping session, we acquired an image of the superficial brain vasculature pattern under the same field of view. We acquired a similar brain vasculature image under the large field of view two-photon microscope. These two reference images were manually aligned and used to directly locate V1 and PPC locations for two-photon imaging. The location for RSC imaging was positioned adjacent to the midline and about 300 μm anterior of the PPC location.

### General analyses

Statistical estimates and significance were generally generated with hierarchical bootstrapping (Saravanan et al., 2020), and data are reported as mean ± sem of hierarchical bootstrap distributions, unless noted otherwise. Standard error of the mean was calculated as the standard deviation of the means from a bootstrap distribution (n = 10000 resampled datasets). For analyses of optogenetic inhibition effects, resampled datasets were generated by sampling with replacement first at the level of sessions pooled across mice and then at the level of trials. For analyses of calcium imaging data, resampled datasets were generated by resampling at the level of sessions then neurons. For significance testing of differences between bootstrap distributions, the probability that one was greater or less than the other, whichever was smaller, was computed. To obtain a p-value for a two-tailed test with α = 0.05, this probability was doubled. Analyses were performed with custom code in MATLAB. No statistical methods were used to predetermine sample sizes, but our sample sizes were similar to ones in previous publications in the field. Allocation of individual mice into experimental groups, i.e. behavioral tasks, was not randomized, and co-housed mice were trained on the same behavioral task and task sequence. Data collection was not performed blind to the experimental groups. Blinding experimenters would have been challenging as experimenters remained present throughout behavioral sessions to ensure the sessions were running smoothly, and many experimental groups were inferable by observing the virtual reality display and rewarded choices over time. Data collection was performed by four different experimentalists. Analyses were also non-blinded but performed by two different experimentalists. A small number of behavioral sessions were excluded from analysis due to low performance of the mouse on control trials. Imaging sessions were excluded in case of noticeable drift after motion correction.

### Analysis of photoinhibition experiments

#### Effects of photoinhibition on performance

Performance was quantified as ‘fraction correct’, the fraction of trials in which the mouse made the correct choice. Chance performance was 50% correct. Effects of cortical inhibition were measured as ΔFraction Correct, the fraction correct with inhibition minus the fraction correct with the laser steered to the control (off-cortex) spot. Fraction correct and ΔFraction Correct were calculated on a session-basis. For comparisons of control performance and performance with various cortical inhibition targets within a task, significance levels were adjusted with the Bonferroni method.

#### Quantification of learning times

To compare the number of training sessions necessary to achieve expert performance across tasks (Figure 1), training sessions were counted from the first day on the Y-maze, after training on the linear maze, until both of the following performance criteria were reached per session: maximum of 20% “visually guided trials” and average fraction correct of at least 70% correct (switching task) or 85% (simple task and delay task). Note that in the switching task, these performance criteria included all trials per session, including trials following rule switches. For the delay task, an additional performance criterion was a delay length of 50 cm, and only mice without prior photoinhibition sessions in the simple task were included (5 out of 7 delay task mice).

#### Photoinhibition effects on biases and running parameters

Choice biases were calculated per mouse and task (Figure 1—figure supplement 2). For each session, a signed choice bias value for each inhibition target was calculated as: (Frac Corr_left_ – Frac Corr_right_) / (Frac Corr_left_ + Frac Corr_right_). Thus, a signed choice bias of 1 or -1 indicates that the mouse only made left or right choices, respectively. Inhibition effects on running parameters for each inhibition target were calculated by averaging the treadmill velocity for forward running (pitch axis) per trial during inhibition and normalizing each value by the average treadmill pitch velocity in control trials of the same session. The resulting values were averaged within each mouse before averaging across mice.

### Analysis of calcium imaging experiments

#### Pre-processing of imaging data

To correct for motion artifacts, custom code was used as described in detail previously (Chettih & Harvey, 2019): https://github.com/HarveyLab/Acquisition2P_class/tree/motionCorrection. In brief, motion correction was implemented as a sum of shifts on three distinct temporal scales: sub-frame, full-frame, and minutes-to-hour-long warping. After motion correction, regions of interest (ROIs) were extracted with Suite2P (Pachitariu et al., 2016). Afterwards, somatic sources were identified with a custom two-layer convolutional network in MATLAB trained on manually annotated labels to classify ROIs as neural somata, processes, or other (Chettih & Harvey, 2019). Only somatic sources were used. After identifying individual neurons, average fluorescence in each ROI was computed and converted into a normalized change in fluorescence (ΔF/F). We corrected the numerator of the ΔF/F calculation for neuropil by subtracting a scaled version of the neuropil signal estimated per neuron during source extraction:

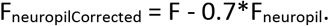

The baseline fluorescence of this trace was estimated as the 8th percentile of fluorescence within a 60 s window (baseline_neuropilCorrected_), and subtracted to get the numerator:

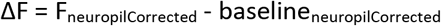

We divided this by the baseline (again 8th percentile of 60 s window) of the raw fluorescence signal to get ΔF/F. The ΔF/F trace per neuron was deconvolved using the constrained AR-1 OASIS method (Friedrich et al., 2017). Decay constants were initialized at two seconds and optimized separately for each neuron.

All analyses were performed on deconvolved activity that was spatially binned along the long axis of the maze (5 cm bins). To be able to compare neural activity across tasks, only correct trials from high performance periods were included (minimum of 80% correct in a window of 10 trials, which excludes periods after rule switches in the switching task). In the switching task, only trials from a single rule (Rule A, i.e. the vertical grating cue/horizontal grating cue requires a right/left choice) were included. Furthermore, for comparisons of trial-type selectivity, noise correlations, or trial-type decoding across tasks, trials were subsampled to the low number of trials per trial type (i.e. horizontal cue / left trial versus vertical cue / right trial) per session in the switching task when considering only high performance trials for Rule A (n = 30 trials per trial type).

#### Trial-type selectivity

To quantify if activity of single neurons was informative about the current trial type, the area under the receiver operating characteristics curve (auROC) was calculated for each bin and averaged per maze segment (first half of stem, second half of stem, Y-arms). Trial-type selectivity was defined as the unsigned version of the auROC: 2*|auROC – 0.5| (Najafi et al., 2020). To identify neurons with significant trial-type selectivity, for each neuron, unsigned auROC values were recomputed 100 times with shuffled trial labels, and the original value was compared to the resulting distribution. Trial-type selectivity was considered significant if the probability of drawing this value from the shuffled distribution was less than 0.01. The fraction of trial-type selective neurons was calculated for each spatial bin and subsequently averaged per maze segment.

#### Trial-type decoding

For each session and area, at each spatial bin, a linear SVM was trained to predict the current trial type (i.e. horizontal cue / left trial versus vertical cue / right trial) using the activity of a subsample of neurons (n = 5, 10, 25, 50, 75, 150 or 200 neurons, activity of each neuron z-scored), with 10-fold cross-validation. This procedure was repeated 40 times for populations of 5 or 10 neurons, and 20 times for populations of 25-200 neurons. For each repetition, the decoding accuracy per bin was calculated as the fraction of test trials in which the trial type was predicted correctly. Decoding accuracy was averaged across spatial bins and repetitions per subsampled population per session. To compare trial-type decoding across tasks, a two-way ANOVA with factors for task and population size was used.

#### Noise correlations

To measure pairwise noise correlations, we calculated the Pearson correlation coefficient for pairs of neurons separately for each trial type, and then averaged the coefficients across trial types. To compare noise correlations across tasks, a two-way ANOVA with factors for task and brain area combination was used. To assess the effect of noise correlations on population information, we disrupted noise correlations by shuffling the order of trials for each neuron independently for each trial type and repeated the trial-type decoding analysis above. We then calculated the difference in decoding accuracy, subtracting the mean accuracy with disrupted noise correlations from the mean accuracy with intact noise correlations, for a given population size and task.

#### Choice decoding based on running parameters

To quantify how well a mouse’s reported choice could be decoded from its running parameters in a given task, a generalized linear model was fit using as predictors the instantaneous treadmill velocities for all axes (pitch, roll, yaw), and the lateral maze position. Running parameters were spatially binned along the maze’s long axis (5 cm bins), and a different model was trained for each bin with 10-fold cross-validation. In photoinhibition experiments, only control trials were used. Only correct trials from high performance periods were used (minimum of 80% correct in a window of 10 trials, which excludes periods after rule switches in the switching task), and in each session, trials were subsampled to the low number of trials per trial type in the switching task when considering only high-performance trials for a single rule per session (n = 30 trials per trial-type). To sub-select calcium imaging sessions with similar running patterns to control for differences in running patterns across tasks (Figure 4—figure supplement 2), we used a session-wise criterion of average choice decoding accuracy of 85-95% in the maze stem.

---

**Figure 1—figure supplement 1.**
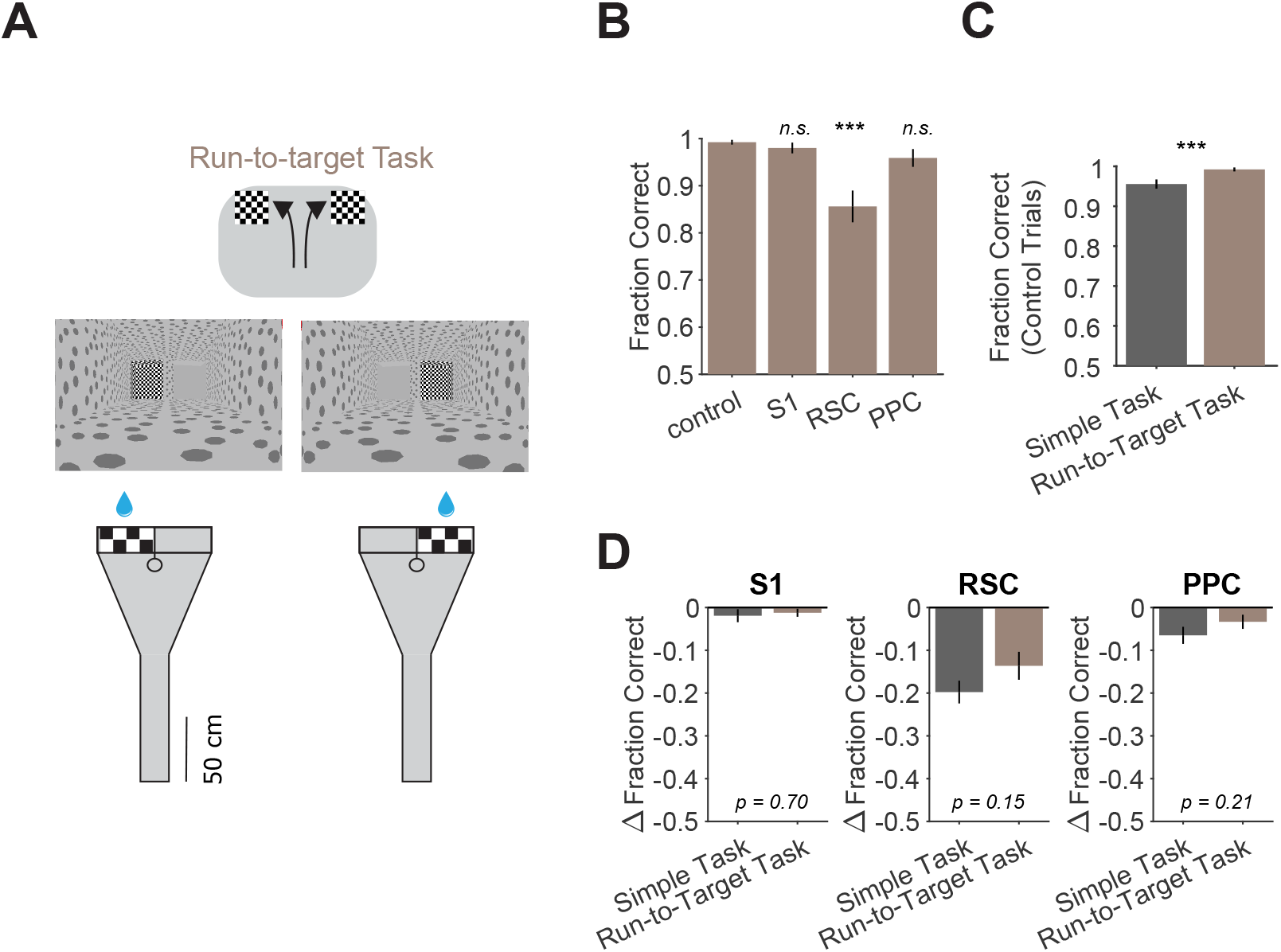
Similar deficits from inhibition in a run-to-target task as in the simple task. **(A)** Schematic and virtual reality screenshots of run-to-target task showing left and right trials. **(B)** Performance in the run-to-target task for each inhibited location across 15 sessions from 3 mice. Bars indicate mean ± sem of a bootstrap distribution of the mean. S1 p = 0.85; RSC p < 0.001; PPC p = 0.16; from bootstrapped distributions of ΔFraction Correct (difference from control performance) compared to 0, two-tailed test, α = 0.05 plus Bonferroni correction. Sessions per mouse: 5 ± 2. Trials per session: 93 ± 11 (control), 26 ± 5 (S1), 24 ± 5 (RSC), 28 ± 6 (PPC), mean ± SD. **(C)** Comparison of performance on control trials in the simple task (same dataset as in Figure 1K) versus the run-to-target task using only the first two laser-on blocks in each session. Bars indicate mean ± sem of a bootstrap distribution of the mean; p < 0.001, two-tailed comparison of bootstrapped Fraction Correct distributions, α = 0.05. Trials per session: 51 ± 23 (simple task), 93 ± 11 (run-to-target task), mean ± SD. **(D)** Comparison of inhibition effects (ΔFraction Correct) in the simple task (same dataset as in Figure 1F) and the run-to-target task for each cortical inhibition location. Bars indicate mean ± sem of a bootstrap distribution of the mean; two-tailed comparisons of bootstrapped ΔFraction Correct distributions, α = 0.05.

**Figure 1—figure supplement 2.**
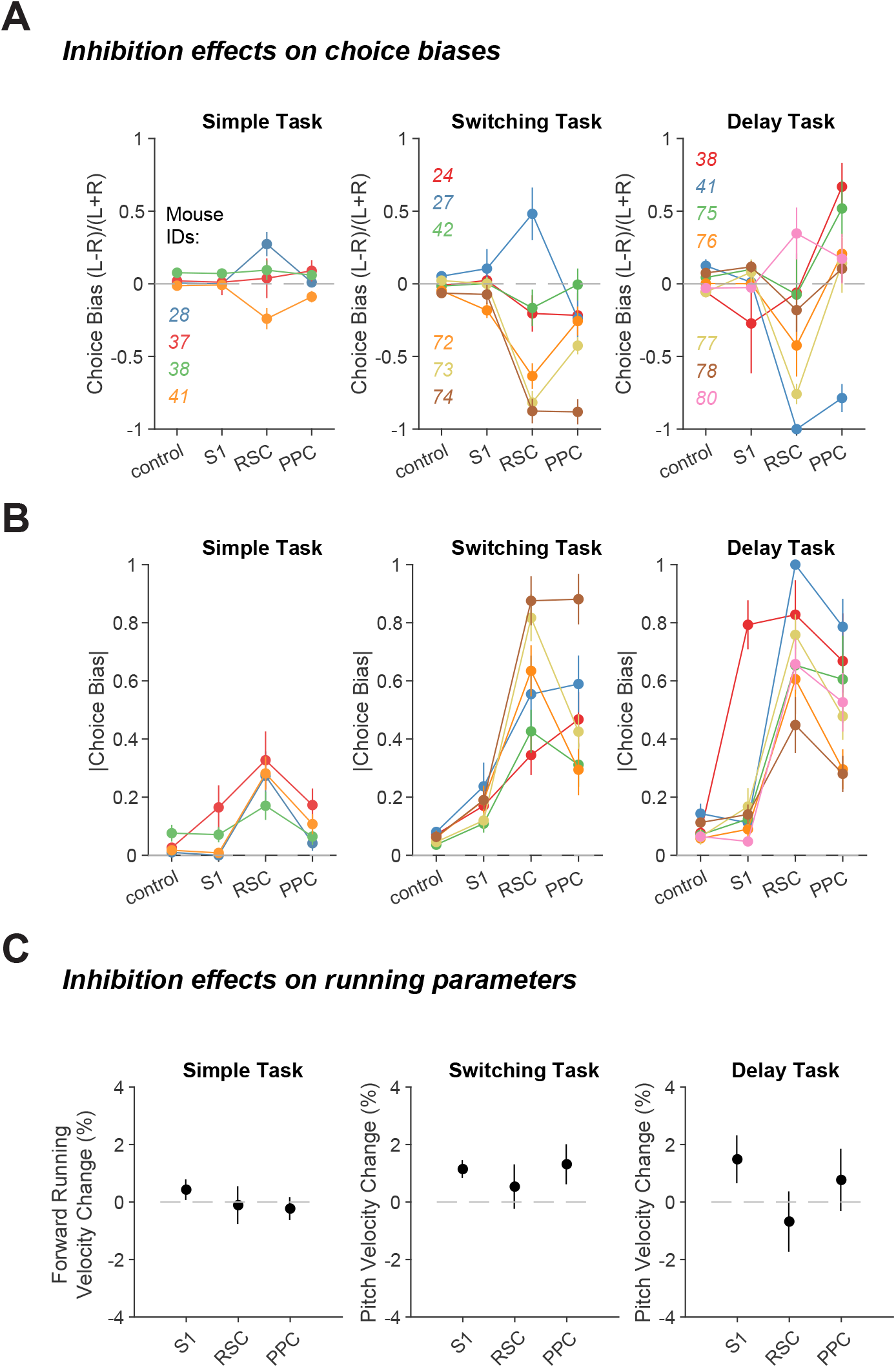
Inhibition effects on choice biases and running parameters across tasks. **(A)** Signed choice bias for each inhibited location in each task. Line colors indicate different mice. Error bars indicate mean ± sem across sessions. Sessions per task per mouse: 11 ± 2 (simple), 15 ± 5 (switching), 9 ± 4 (delay). **(B)** Same as in (**A**), except for the absolute value of choice bias. **(C)** Percent change in forward running velocity with inhibition, averaged across the entire maze. Error bars indicate mean ± sem across mice. Mice per task: 4 (simple), 6 (switching), 7 (delay).

**Figure 1—figure supplement 3.**
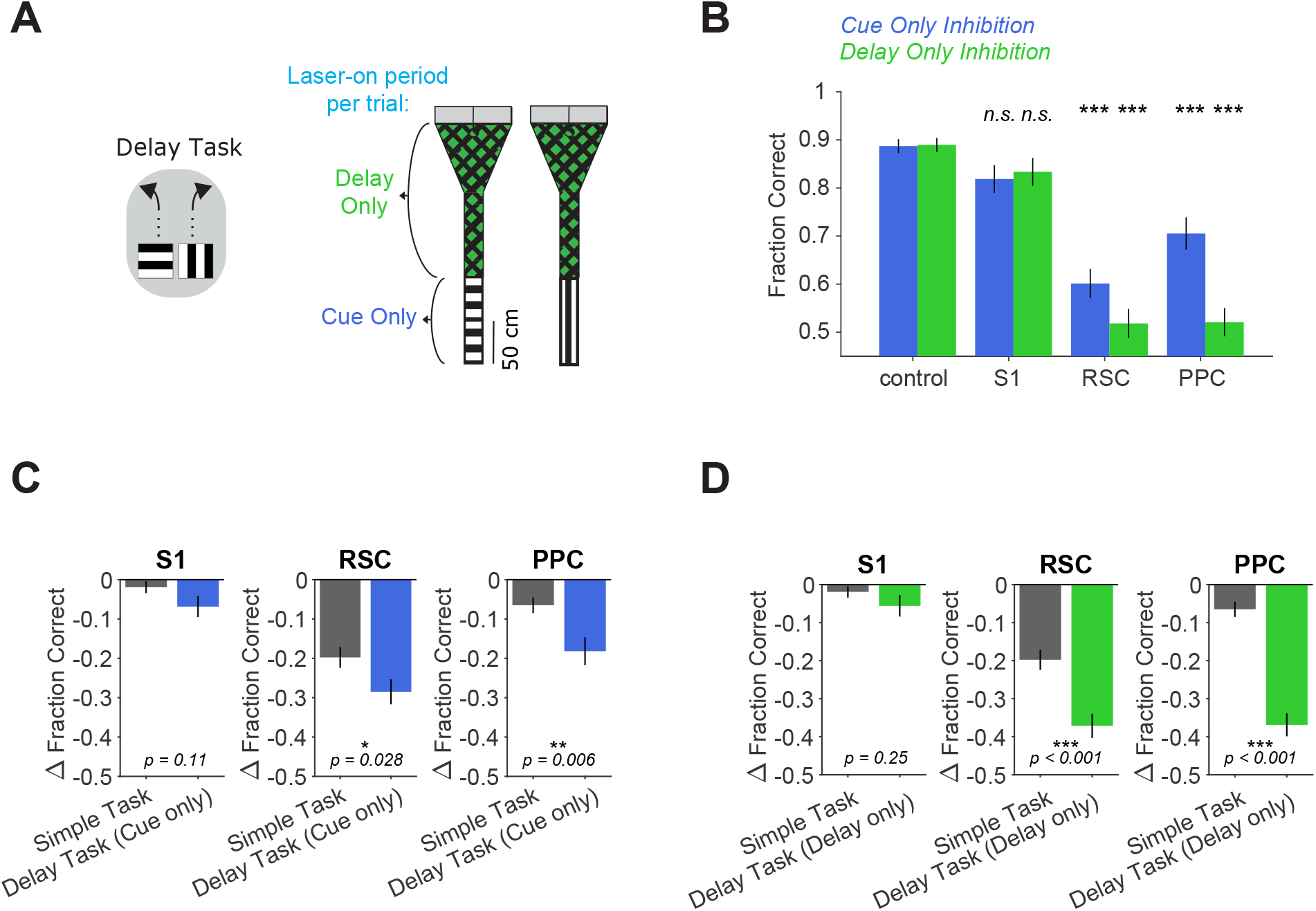
Cue only or delay only inhibition in the delay task. **(A)** Left: schematic of the delay task. Right: Inhibition was restricted to either the cue period only or the delay period only in a given session. **(B)** Performance in the delay task with cue only (blue, 48 sessions from 5 mice) or delay only (green, 45 sessions from 5 mice) inhibition for each inhibited location. Bars indicate mean ± sem of a bootstrap distribution of the mean. For cue only or delay only inhibition individually, inhibition performance was compared to control performance, two-tailed test, α = 0.05 plus Bonferroni correction. Cue only: S1 p = 0.09; RSC p < 0.001; PPC p < 0.001. Sessions per mouse: 10 ± 2. Trials per session: 59 ± 16 (control), 14 ± 6 (S1), 14 ± 5 (RSC), 15 ± 5 (PPC), mean ± SD. Delay only: S1 p = 0.27; RSC p < 0.001; PPC p < 0.001. Sessions per mouse: 9 ± 2. Trials per session: 61 ± 15 (control), 14 ± 5 (S1), 15 ± 4 (RSC), 15 ± 5 (PPC), mean ± SD. **(C)** Comparison of inhibition effects (ΔFraction Correct) in the simple task (same dataset as in Figure 1F) and the delay task with cue inhibition only for each cortical location. Bars indicate mean ± sem of a bootstrap distribution of the mean; two-tailed comparisons of bootstrapped ΔFraction Correct distributions, α = 0.05. **(D)** Similar to **(C)**, but for delay inhibition only in the delay task.

**Figure 1—figure supplement 4.**
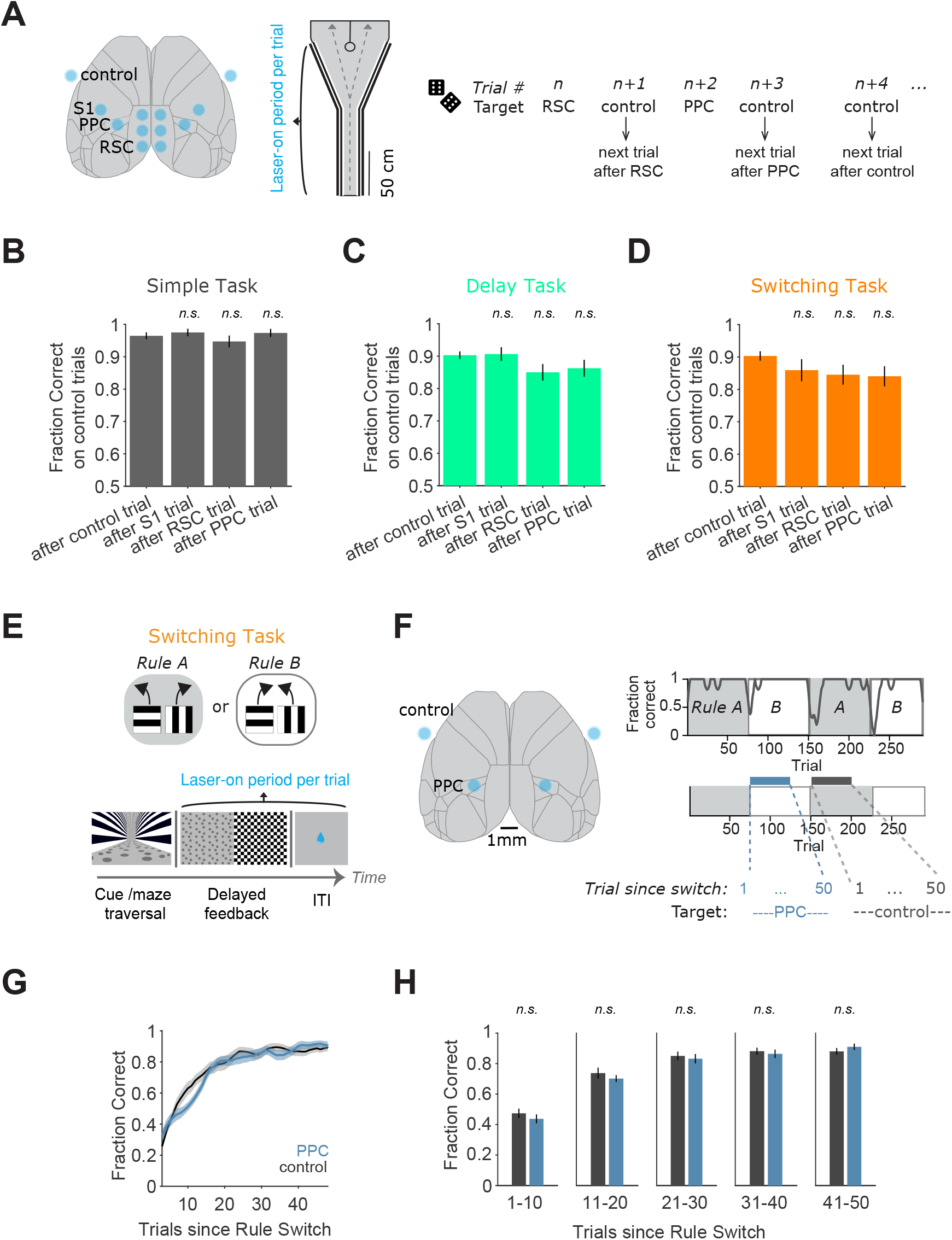
Performance on trials following inhibition and rule switching with inhibition during the feedback/ITI period. **(A)** Left: Schematic of the inhibition locations (same as in Figure 1). Middle: inhibition lasted from trial onset throughout maze traversal. Right: As in Figure 1, inhibition target locations per trial were randomly interleaved. Analysis here used performance on control trials that directly followed inhibition of the labeled location on the preceding trial. **(B)** Performance on control trials immediately following an inhibition trial, for the simple task, for each inhibited location across 45 sessions from 4 mice. Bars indicate mean ± sem of a bootstrap distribution of the mean. S1 p = 1; RSC p = 1; PPC p = 1; from bootstrapped distributions of ΔFraction Correct (difference from control performance) compared to 0, two-tailed test, α = 0.05 plus Bonferroni correction. Sessions per mouse: 11 ± 2. Trials per session: 22 ± 12 (control), 8 ± 3 (S1), 8 ± 4 (RSC), 9 ± 4 (PPC), mean ± SD. **(C)** Similar to **(B)**, except for the delay task. 62 sessions from 7 mice. S1 p = 1; RSC p = 0.12; PPC p = 0.50. Sessions per mouse: 9 ± 4. Trials per session: 29 ± 8 (control), 8 ± 3 (S1), 9 ± 4 (RSC), 7 ± 3 (PPC). **(D)** Similar to **(B)**, except for the switching task (Rule A trials only). 89 sessions from 6 mice. S1 p = 0.66; RSC p = 0.27; PPC p = 0.19. Sessions per mouse: 15 ± 5. Trials per session: 13 ± 6 (control), 4 ± 2 (S1), 5 ± 2 (RSC), 4 ± 2 (PPC). **(E)** Top: Schematic of the switching task. Bottom: schematic of a single trial with inhibition during the feedback and ITI period. **(F)** Left: Schematic of PPC and control targets. Right top: Example behavioral performance in one session in the switching task. Right bottom: inhibition blocks of 50 trials started after a rule switch, with inhibition during the feedback/ITI period. The same area was targeted on every trial. **(G)** Average performance after a rule switch with PPC (blue) or control (black) inhibition on every trial during the feedback/ITI. 33 sessions from 4 mice (8 ± 2 sessions per mouse, mean ± SD). Shading indicates mean ± sem across sessions. Thin lines indicate single sessions. Fraction Correct was Gaussian-filtered (window of 7 trials, sigma of 3 trials) and smoothed again with a moving average filter of 3 trials for plotting. **(H)** Comparison of mean performance with PPC versus control inhibition after a rule switch in bins of 10 trials. Error bars indicate mean ± SEM across sessions, gray lines show single sessions (n = 33). Paired two-sided t-tests. p (trials 1-10): 0.42; p (trials 11-20): 0.43; p (trials 21-30): 0.64; p (trials 31-40): 0.66; p (trials 41-50): 0.34.

**Figure 3—figure supplement 1.**
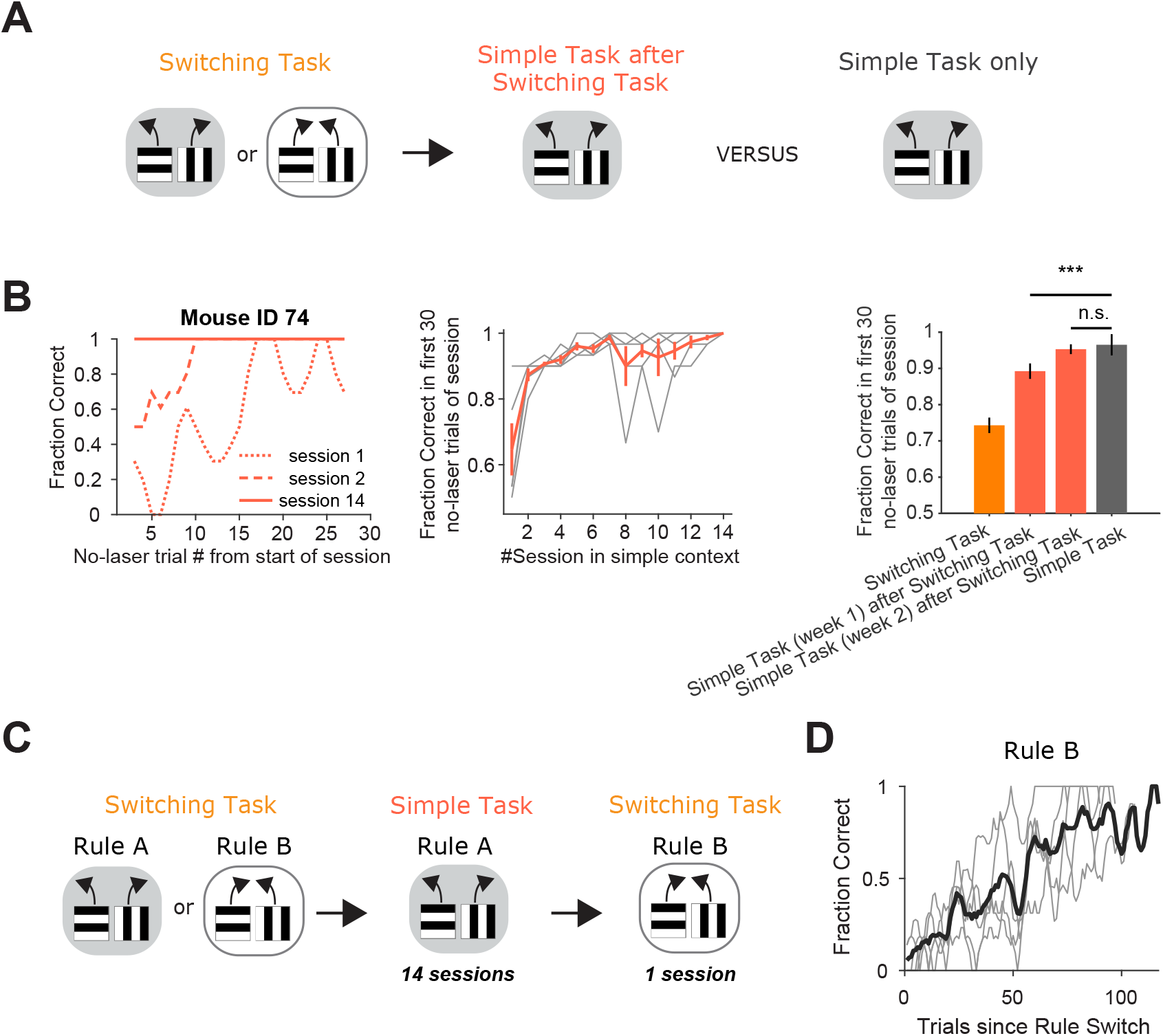
Mice with experience in the switching task perform at high levels on no-laser trials in the simple task and can still switch rules. **(A)** Schematic of the training history sequence. One group of mice was trained on the switching task and then permanently transitioned to the simple task. This group of mice was compared to another group trained only on the simple task. **(B)** Left: Performance for an example mouse with switching task experience across the first 30 trials per session for sessions 1, 2, and 14 in the simple task. Each session started with a block of no-laser trials (minimum of 50 trials) before the first laser-on block. Fraction Correct was Gaussian-smoothed with a window size of 7 trials, sigma of 3 trials. Middle: Average performance in the first 30 no-laser trials of a session in the simple task. Gray lines: individual mice. Bars indicate mean ± sem across mice (n = 5). Right: Average initial performance in the switching task (orange, 90 sessions from 6 mice), in each week in the simple task of mice with previous switching task experience (red, 35 and 34 sessions from 5 mice in week 1 and 2, respectively), and in the simple task in mice with simple-task-only experience (gray, 45 sessions from 4 mice). Bars indicate mean ± sem across sessions. Unpaired two-sided t-tests comparing performance in the simple task with and without switching task experience. Week 1: p = 0.00096, week 2: p = 0.43. **(C)** Schematic of task sequence: After training in the switching task, mice were transitioned to the simple task without any rule switches for 14 days. Then for a single session, mice were exposed to the rule they had not been exposed to for 14 days. **(D)** Performance in the rule mice had not experienced for 14 days. Gray lines: individual mice. Black line: average across mice. Lines are only shown until the next rule switch, which different mice encountered at different numbers of trials based on their performance.

**Figure 3—figure supplement 2.**
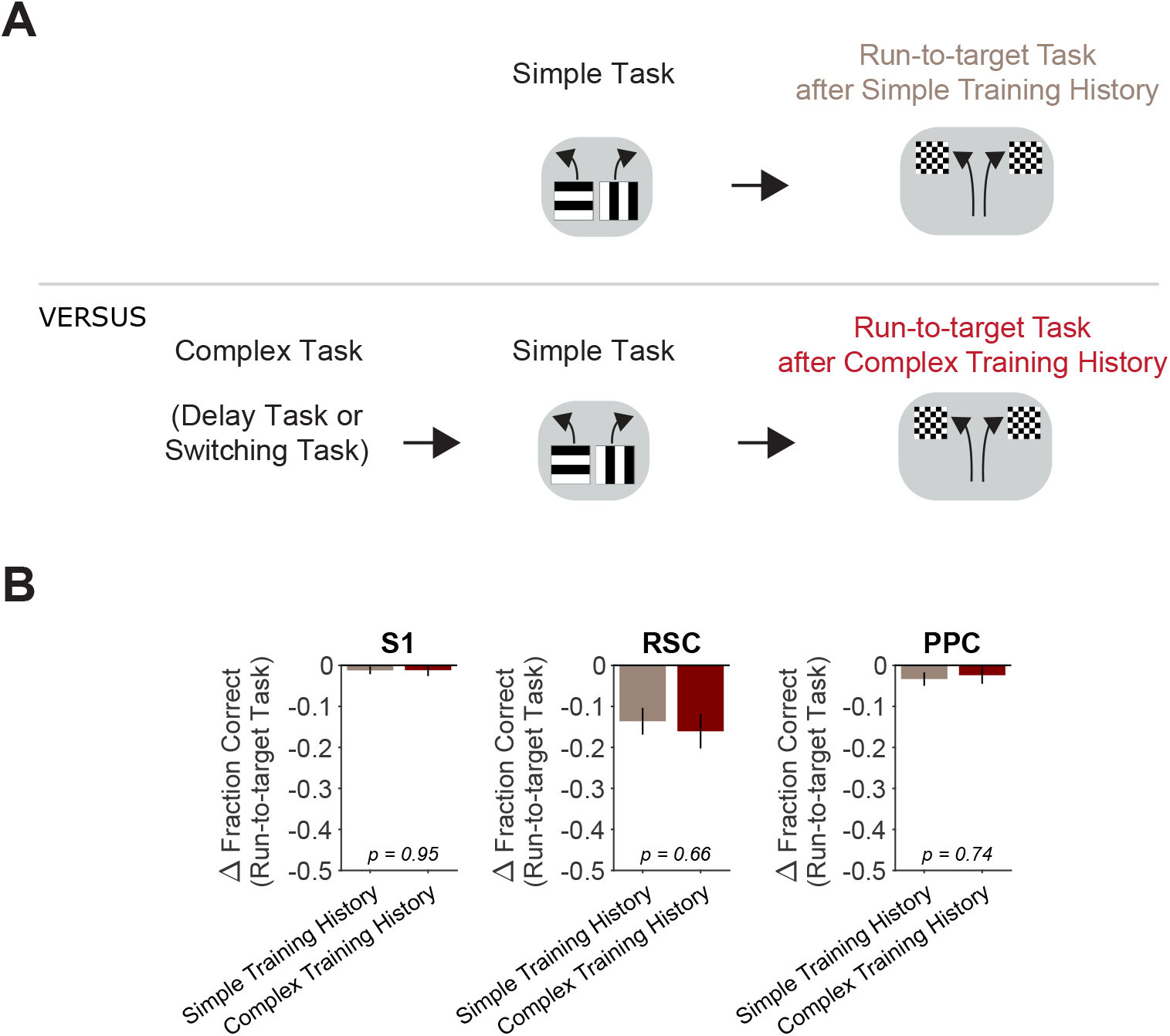
Increased cortical involvement in the simple task after complex task experience does not generalize to the run-to-target task. **(A)** Schematic of task training sequence. One group of mice was trained in the simple task and then transitioned to the run-to-target task. Another group of mice was first trained in a complex task (switching or delay task), then transitioned to the simple task for 14 days, and then tested on the run-to-target task. **(B)** Comparison of inhibition effects (ΔFraction Correct) in the run-to-target task in mice with simple task-only versus complex and simple task experience, for each cortical inhibition location. Bars indicate mean ± sem of a bootstrap distribution of the mean; two-tailed comparisons of bootstrapped ΔFraction Correct distributions, α = 0.05. Simple task-only experience (same dataset as in Figure 1— figure supplement 1): 15 sessions from 3 mice, 5 ± 2 sessions per mouse. Trials per session: 93 ± 11 (control), 26 ± 5 (S1), 24 ± 5 (RSC), 28 ± 6 (PPC), mean ± SD. Complex task and simple task experience: 11 sessions from 3 mice, 4 ± 2 sessions per mouse. Trials per session: 94 ± 4 (control), 28 ± 2 (S1), 23 ± 2 (RSC), 28 ± 4 (PPC), mean ± SD.

**Figure 4—figure supplement 1.**
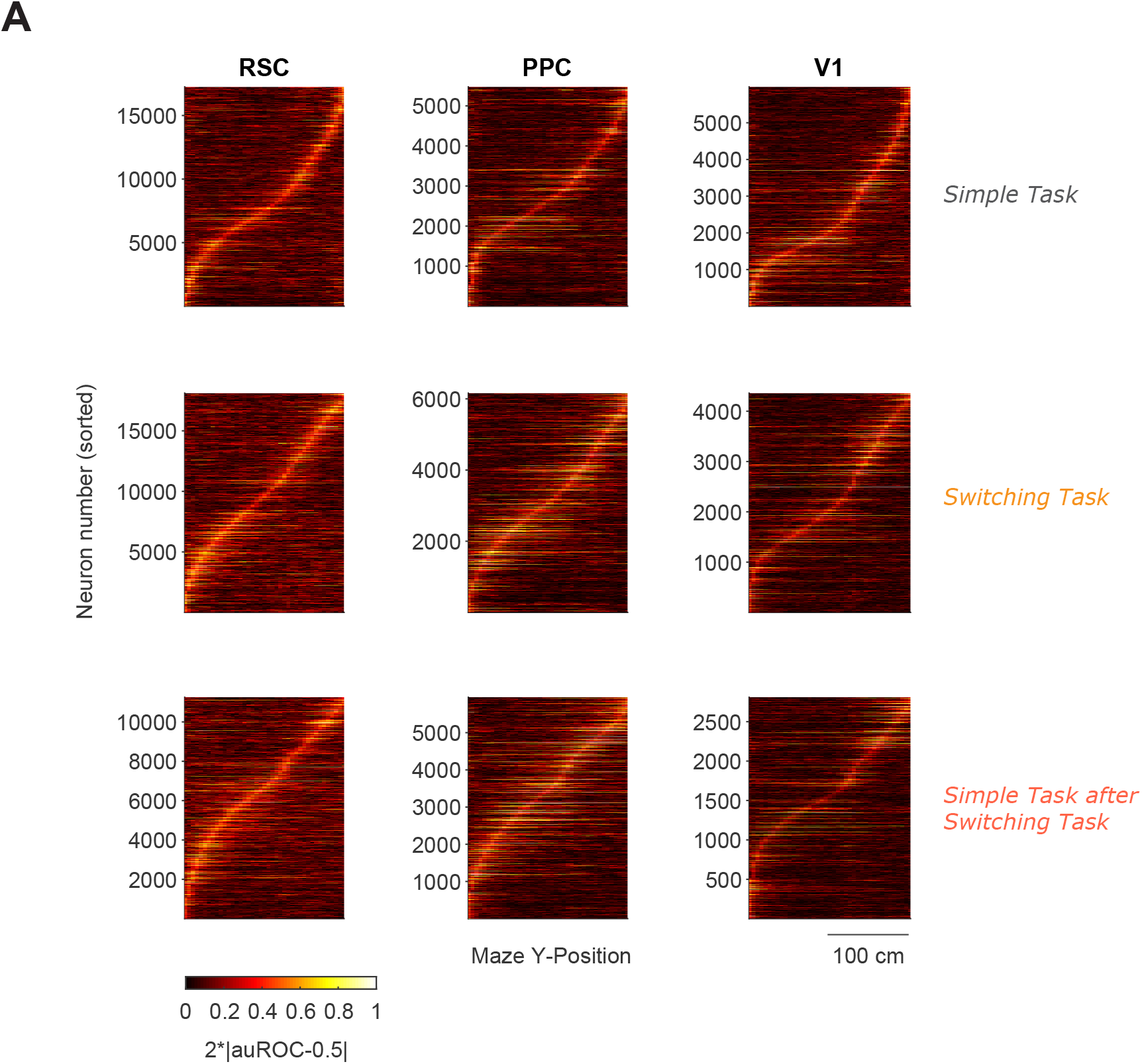
Neuronal trial-type selectivity is sequentially organized. **(A)** Trial-type selectivity (absolute deviation of auROC from chance level, 2*|auROC-0.5|) was calculated per neuron and spatial maze bin (5 cm bin size) and is shown for all neurons pooled across mice and sessions in each cortical area and task. Neurons were sorted by the maze position of their selectivity peak. Mice per task: simple (3), switching (3), simple after switching (2).

**Figure 4—figure supplement 2.**
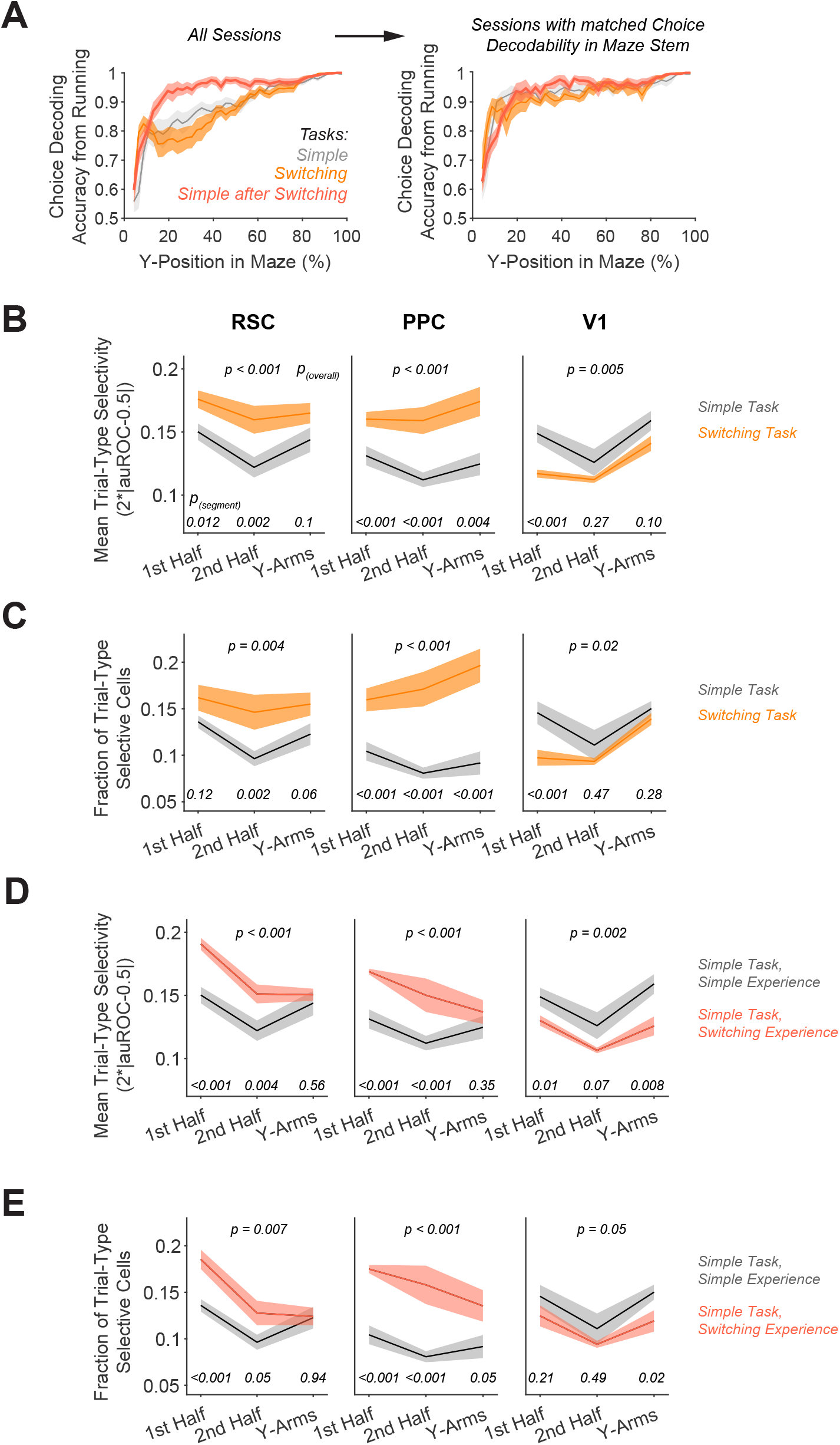
Trial-type selectivity for sessions with similar running patterns. **(A)** Left: Decoding accuracy of the reported choice using instantaneous treadmill velocities and lateral position, binned along the maze’s long axis (5 cm bins). Shading indicates mean ± sem across all sessions per task. Sessions per task: 12 (simple), 12 (switching), 8 (simple after switching). Right: Decoding accuracy using a subset of sessions with similar running patterns, i.e. with average choice decoding accuracy in the maze stem ranging from 85-95%. Sessions per task: 4 (simple), 5 (switching), 5 (simple after switching). **(B)** For sessions with similar running patterns, in each area, mean trial-type selectivity levels across cells by maze segment are compared in the simple versus the switching task (Rule A trials only). Shading indicates mean ± sem of bootstrapped distributions. p_(segment)_ shows p values of two-tailed comparisons of bootstrapped distributions per maze segment. p_(overall)_ shows the p value for the task factor from a two-way ANOVA (factors: task and maze segment). **(C)** For sessions with similar running patterns, similar to **(B)**, except for the fraction of trial-type selective cells as determined from comparing each cell’s selectivity value to a distribution with shuffled trial labels (significance threshold of p < 0.01). **(D)** For sessions with similar running patterns, similar to **(B)**, but for the simple task after switching task experience. **(E)** For sessions with similar running patterns, similar to **(C)**, but for the simple task after switching task experience.

## References

Akrami, A., Kopec, C. D., Diamond, M. E., & Brody, C. D. (2018). Posterior parietal cortex represents sensory history and mediates its effects on behavior. Nature, 182246. https://doi.org/10.1101/182246

Alexander, A. S., & Nitz, D. a. (2015). Retrosplenial cortex maps the conjunction of internal and external spaces. Nature Neuroscience, 18(July), 1–12. https://doi.org/10.1038/nn.4058

Aronov, D., & Tank, D. W. (2014). Engagement of Neural Circuits Underlying 2D Spatial Navigation in a Rodent Virtual Reality System. Neuron, 84(2), 442–456. https://doi.org/10.1016/j.neuron.2014.08.042

Averbeck, B. B., Latham, P. E., & Pouget, A. (2006). Neural correlations, population coding and computation. Nature Reviews. Neuroscience, 7(5), 358–366. https://doi.org/10.1038/nrn1888

Buschman, T. J., Siegel, M., Roy, J. E., & Miller, E. K. (2011). Neural substrates of cognitive capacity limitations. Proceedings of the National Academy of Sciences of the United States of America, 108(27), 11252–11255. https://doi.org/10.1073/pnas.1104666108

Ceballo, S., Piwkowska, Z., Bourg, J., Daret, A., & Bathellier, B. (2019). Targeted Cortical Manipulation of Auditory Perception. Neuron, 1–12. https://doi.org/10.1016/j.neuron.2019.09.043

Chettih, S. N., & Harvey, C. D. (2019). Single-neuron perturbations reveal feature-specific competition in V1. Nature, 567(7748), 334–340. https://doi.org/10.1038/s41586-019-0997-6

Chowdhury, S. A., & DeAngelis, G. C. (2008). Fine Discrimination Training Alters the Causal Contribution of Macaque Area MT to Depth Perception. Neuron, 60(2), 367–377. https://doi.org/10.1016/j.neuron.2008.08.023

Cohen, M. R., & Kohn, A. (2011). Measuring and interpreting neuronal correlations. Nature Neuroscience, 14(7), 811–819. https://doi.org/10.1038/nn.2842

Driscoll, L. N., Pettit, N. L., Minderer, M., Chettih, S. N., & Harvey, C. D. (2017). Dynamic Reorganization of Neuronal Activity Patterns in Parietal Cortex. Cell, 170(5), 986–999.e16. https://doi.org/10.1016/j.cell.2017.07.021

Erlich, J. C., Bialek, M., & Brody, C. D. (2011). A cortical substrate for memory-guided orienting in the rat. Neuron, 72(2), 330–343. https://doi.org/10.1016/j.neuron.2011.07.010

Fischer, L. F., Mojica Soto-Albors, R., Buck, F., & Harnett, M. T. (2020). Representation of visual landmarks in retrosplenial cortex. ELife, 9, 811430. https://doi.org/10.7554/eLife.51458

Freedman, D. J., & Ibos, G. (2018). An Integrative Framework for Sensory, Motor, and Cognitive Functions of the Posterior Parietal Cortex. Neuron, 97(6), 1219–1234. https://doi.org/10.1016/j.neuron.2018.01.044

Friedrich, J., Zhou, P., & Paninski, L. (2017). Fast online deconvolution of calcium imaging data. PLOS Computational Biology, 13(3), e1005423. https://doi.org/10.1371/journal.pcbi.1005423

Fuster, J. M. (1997). Network memory. Trends in Neurosciences, 20(10), 451–459. https://doi.org/10.1016/S0166-2236(97)01128-4

Goard, M. J., Pho, G. N., Woodson, J., & Sur, M. (2016). Distinct roles of visual, parietal, and frontal motor cortices in memory-guided sensorimotor decisions. ELife, 5(AUGUST), 1–30. https://doi.org/10.7554/eLife.13764

Gold, J. I., & Shadlen, M. N. (2007). The neural basis of decision making. Annu Rev Neurosci, 30, 535– 574. https://doi.org/10.1146/annurev.neuro.29.051605.113038

Golub, M. D., Sadtler, P. T., Oby, E. R., Quick, K. M., Ryu, S. I., Tyler-Kabara, E. C., Batista, A. P., Chase, S. M., & Yu, B. M. (2018). Learning by neural reassociation. Nature Neuroscience, 21(4), 607–616. https://doi.org/10.1038/s41593-018-0095-3

Guo, Z., Li, N., Huber, D., Ophir, E., Gutnisky, D., Ting, J., Feng, G., & Svoboda, K. (2014). Flow of cortical activity underlying a tactile decision in mice. Neuron, 81(1), 179–194. https://doi.org/10.1016/j.neuron.2013.10.020

Hanks, T. D., Ditterich, J., & Shadlen, M. N. (2006). Microstimulation of macaque area LIP affects decision-making in a motion discrimination task. Nature Neuroscience, 9(5), 682–689. https://doi.org/10.1038/nn1683

Harvey, C. D., Coen, P., & Tank, D. W. (2012). Choice-specific sequences in parietal cortex during a virtual-navigation decision task. Nature, 484(7392), 62–68. https://doi.org/10.1038/nature10918

Harvey, C. D., Collman, F., Dombeck, D. A., & Tank, D. W. (2009). Intracellular dynamics of hippocampal place cells during virtual navigation. Nature, 461(7266), 941–946. https://doi.org/10.1038/nature08499

Hwang, E. J., Dahlen, J. E., Hu, Y. Y., Aguilar, K., Yu, B., Mukundan, M., Mitani, A., & Komiyama, T. (2019). Disengagement of motor cortex from movement control during long-term learning. Science Advances, 5(10), 1–13. https://doi.org/10.1126/sciadv.aay0001

Inagaki, H. K., Inagaki, M., Romani, S., & Svoboda, K. (2018). Low-dimensional and monotonic preparatory activity in mouse anterior lateral motor cortex. Journal of Neuroscience, 38(17), 4163– 4185. https://doi.org/10.1523/JNEUROSCI.3152-17.2018

Kafashan, M. M., Jaffe, A. W., Chettih, S. N., Nogueira, R., Arandia-Romero, I., Harvey, C. D., Moreno-Bote, R., & Drugowitsch, J. (2021). Scaling of sensory information in large neural populations shows signatures of information-limiting correlations. Nature Communications, 12(1), 1–16. https://doi.org/10.1038/s41467-020-20722-y

Kalatsky, V. A., & Stryker, M. P. (2003). New paradigm for optical imaging: Temporally encoded maps of intrinsic signal. Neuron, 38(4), 529–545. https://doi.org/10.1016/S0896-6273(03)00286-1

Katz, L., Yates, J., Pillow, J. W., & Huk, A. (2016). Dissociated functional significance of choice-related activity across the primate dorsal stream. Nature, 535(7611), Salt Lake City USA. https://doi.org/10.1038/nature18617

Kawai, R., Markman, T., Poddar, R., Ko, R., Fantana, A. L., Dhawale, A. K., Kampff, A. R., & Ölveczky, B. P. (2015). Motor Cortex Is Required for Learning but Not for Executing a Motor Skill. Neuron, 86(3), 800–812. https://doi.org/10.1016/j.neuron.2015.03.024

Kim, T. H., Zhang, Y., R^ Ome Lecoq, J., Zeng, H., Niell, C. M., Schnitzer Correspondence, M. J., Jung, J. C., Li, J., & Schnitzer, M. J. (2016). Long-Term Optical Access to an Estimated One Million Neurons in the Live Mouse Cortex. Cell Reports, 17, 3385–3394. https://doi.org/10.1016/j.celrep.2016.12.004

Kılıç, K., Desjardins, M., Tang, J., Thunemann, M., Sunil, S., Erdener, Ş. E., Postnov, D. D., Boas, D. A., & Devor, A. (2021). Chronic Cranial Windows for Long Term Multimodal Neurovascular Imaging in Mice. Frontiers in Physiology, 11(January), 1–10. https://doi.org/10.3389/fphys.2020.612678

Koay, S. A., Thiberge, S. Y., Brody, C. D., & Tank, D. W. (2020). Amplitude modulations of cortical sensory responses in pulsatile evidence accumulation. ELife, 9, 1–49. https://doi.org/10.7554/eLife.60628

Lashley, K. S. (1931). Mass Action in Cerebral Function. Science, 73(1888), 245–254. https://doi.org/10.1126/science.73.1888.245

Li, N., Chen, S., Guo, Z. V, Chen, H., Huo, Y., Inagaki, H. K., Chen, G., Davis, C., Hansel, D., Guo, C., & Svoboda, K. (2019). Spatiotemporal constraints on optogenetic inactivation in cortical circuits. ELife, 8. https://doi.org/10.7554/eLife.48622

Licata, A. M., Kaufman, M. T., Raposo, D., Ryan, M. B., Sheppard, J. P., & Churchland, A. K. (2017). Posterior parietal cortex guides visual decisions in rats. Journal of Neuroscience, 37(19), 4954– 4966. https://doi.org/10.1523/JNEUROSCI.0105-17.2017

Liu, L. D., & Pack, C. C. (2017). The Contribution of Area MT to Visual Motion Perception Depends on Training. Neuron, 95(2), 436–446.e3. https://doi.org/10.1016/j.neuron.2017.06.024

Lyamzin, D., & Benucci, A. (2019). The mouse posterior parietal cortex: Anatomy and functions. Neuroscience Research, 140, 14–22. https://doi.org/10.1016/j.neures.2018.10.008

Marshel, J. H., Garrett, M. E., Nauhaus, I., & Callaway, E. M. (2011). Functional specialization of seven mouse visual cortical areas. Neuron, 72(6), 1040–1054. https://doi.org/10.1016/j.neuron.2011.12.004

Minderer, M., Brown, K. D., & Harvey, C. D. (2019). The Spatial Structure of Neural Encoding in Mouse Posterior Cortex during Navigation. Neuron, 102(1), 232–248.e11. https://doi.org/10.1016/j.neuron.2019.01.029

Morcos, A. S., & Harvey, C. D. (2016). History-dependent variability in population dynamics during evidence accumulation in cortex. Nature Neuroscience, 19(12), 1672–1681. https://doi.org/10.1038/nn.4403

Najafi, F., Elsayed, G. F., Cao, R., Pnevmatikakis, E., Latham, P. E., Cunningham, J. P., & Churchland, A. K. (2020). Excitatory and Inhibitory Subnetworks Are Equally Selective during Decision-Making and Emerge Simultaneously during Learning. Neuron, 105(1), 165–179.e8. https://doi.org/10.1016/j.neuron.2019.09.045

Oby, E. R., Golub, M. D., Hennig, J. A., Degenhart, A. D., Tyler-Kabara, E. C., Yu, B. M., Chase, S. M., & Batista, A. P. (2019). New neural activity patterns emerge with long-term learning. Proceedings of the National Academy of Sciences of the United States of America, 116(30), 15210–15215. https://doi.org/10.1073/pnas.1820296116

Pachitariu, M., Stringer, C., Schröder, S., Dipoppa, M., Rossi, L. F., Carandini, M., & Harris, K. D. (2016). Suite2p : beyond 10, 000 neurons with standard two-photon microscopy. BioRxiv, 1–14. https://doi.org/10.1101/061507

Panzeri, S., Schultz, S. R., Treves, A., & Rolls, E. T. (1999). Correlations and the encoding of information in the nervous system. Proceedings of the Royal Society B: Biological Sciences, 266(1423), 1001–1012. https://doi.org/10.1098/rspb.1999.0736

Pinto, L., Rajan, K., DePasquale, B., Thiberge, S. Y., Tank, D. W., & Brody, C. D. (2019). Task-Dependent Changes in the Large-Scale Dynamics and Necessity of Cortical Regions. Neuron, 104. https://doi.org/10.1016/j.neuron.2019.08.025

Plitt, M. H., & Giocomo, L. M. (2021). Experience-dependent contextual codes in the hippocampus. Nature Neuroscience, 24(May). https://doi.org/10.1038/s41593-021-00816-6

Raposo, D., Kaufman, M. T., & Churchland, A. K. (2014). A category-free neural population supports evolving demands during decision-making. Nature Neuroscience, 17(12), 1784–1792. https://doi.org/10.1038/nn.3865

Resulaj, A., Ruediger, S., Olsen, S. R., & Scanziani, M. (2018). First spikes in visual cortex enable perceptual discrimination. ELife, 7, e34044. https://doi.org/10.7554/eLife.34044

Ruff, D. A., & Cohen, M. R. (2019). Simultaneous multi-area recordings suggest that attention improves performance by reshaping stimulus representations. Nature Neuroscience, 22(October). https://doi.org/10.1038/s41593-019-0477-1

Sadtler, P. T., Quick, K. M., Golub, M. D., Chase, S. M., Ryu, S. I., Tyler-Kabara, E. C., Yu, B. M., & Batista, A. P. (2014). Neural constraints on learning. Nature, 512(7515), 423–426. https://doi.org/10.1038/nature13665

Salzman, C. D., Britten, K. H., & Newsome, W. T. (1990). Cortical Microstimulation Influences Perceptual Judgements of Motion Direction. Nature, 346(July), 174–177. https://www.nature.com/articles/346174a0.pdf

Saravanan, V., Berman, G. J., & Sober, S. J. (2020). Application of the hierarchical bootstrap to multi-level data in neuroscience. Neuron Behav Data Anal Theory. https://doi.org/10.1101/819334

Sarma, A., Masse, N. Y., Wang, X. J., & Freedman, D. J. (2015). Task-specific versus generalized mnemonic representations in parietal and prefrontal cortices. Nature Neuroscience, 19(1), 143– 149. https://doi.org/10.1038/nn.4168

Sharpe, M. J., Batchelor, H. M., Mueller, L. E., Gardner, M. P. H., & Schoenbaum, G. (2021). Past experience shapes the neural circuits recruited for future learning. Nature Neuroscience, 24(3), 391–400. https://doi.org/10.1038/s41593-020-00791-4

Sofroniew, N. J., Flickinger, D., King, J., & Svoboda, K. (2016). A large field of view two-photon mesoscope with subcellular resolution for in vivo imaging. ELife, 5(JUN2016), 1–20. https://doi.org/10.7554/eLife.14472

Yang, Y., & Zador, A. M. (2012). Differences in sensitivity to neural timing among cortical areas. Journal of Neuroscience, 32(43), 15142–15147. https://doi.org/10.1523/JNEUROSCI.1411-12.2012

Zhang, S., Xu, M., Chang, W.-C., Ma, C., Hoang Do, J. P., Jeong, D., Lei, T., Fan, J. L., & Dan, Y. (2016). Organization of long-range inputs and outputs of frontal cortex for top-down control. Nature Neuroscience, 19(12). https://doi.org/10.1038/nn.4417

Zhou, Y., & Freedman, D. J. (2019). Posterior parietal cortex plays a causal role in perceptual and categorical decisions. Science, 365(July), 180–185. https://doi.org/https://doi.org/10.1126/science.aaw8347

Znamenskiy, P., & Zador, A. M. (2013). Corticostriatal neurons in auditory cortex drive decisions during auditory discrimination. Nature, 497(7450), 482–485. https://doi.org/10.1038/nature12077

Zohary, E., Shadlen, M. N., & Newsome, W. T. (1994). Correlated neuronal discharge rate and its implications for psychophysical performance. Nature, 370(6485), 140–143. https://doi.org/10.1038/370140a0

